# Macrocybin, a mushroom natural triglyceride, reduces tumor growth *in vitro* and *in vivo* through caveolin-mediated interference with the actin cytoskeleton

**DOI:** 10.1101/2020.12.10.418863

**Authors:** Marcos Vilariño, Josune García-Sanmartín, Laura Ochoa-Callejero, Alberto López-Rodríguez, Jaime Blanco-Urgoiti, Alfredo Martínez

## Abstract

Mushrooms have been used for millennia as cancer remedies. Our goal was to screen several species from the rain forest of Costa Rica looking for new antitumor molecules. Mushroom extracts were screened using two human cell lines: A549 (lung adenocarcinoma) and NL20 (immortalized normal lung epithelium). Extracts able to kill tumor cells while preserving nontumor cells were considered “anticancer”. The mushroom with better properties was *Macrocybe titans*. Positive extracts were fractionated further and tested for biological activity on the cell lines. The chemical structure of the active compound was partially elucidated through nuclear magnetic resonance, mass spectrometry, and other ancillary techniques. Chemical analysis showed that the active molecule was a triglyceride containing oleic acid, palmitic acid, and a more complex fatty acid with 2 double bonds. Synthesis of all possible triglycerides and biological testing identified the natural compound, which was named Macrocybin. A xenograft study showed that Macrocybin significantly reduces A549 tumor growth. In addition, Macrocybin treatment resulted in the upregulation of Caveolin-1 expression and the disassembly of the actin cytoskeleton in tumor cells (but not in normal cells). In conclusion, we have shown that Macrocybin constitutes a new biologically active compound that may be taken into consideration for cancer treatment.

## 1. Introduction

Discovery of new anticancer drugs can be accomplished through different approaches, including screening of natural products, testing of synthetic compound libraries, computer-assisted design, and machine learning, among others [(1)]. Living organisms constitute an almost unlimited source of compounds with potential biological activity and, so far, have provided the majority of currently approved therapies for the treatment of cancer [(2, 3)]. A few examples include taxol which was obtained from the bark of the Pacific Yew tree [(4, 5)], vinblastine and related alkaloids from *Vinca* [(6)], camptothecin from the bark of a Chinese tree [(7)], or trabectedin and other drugs which were purified from marine organisms [(8)]. Besides, many natural products are the basis for further chemical development and rational design of synthetic drugs [(9)].

The mechanisms of action by which natural products reduce cancer cell growth are diverse. In the case of taxanes, a direct interaction with tubulin results in cytoskeleton hyperpolymerization and mitosis arrest [(10)]; vinblastine has the reverse effect, depolymerizing the cytoskeleton, but the same final outcome, stopping cell division [(10)]; still others, such as some mushroom extracts, induce immunomodulation [(11, 12)], or affect other cancer hallmarks [(13)].

Mushrooms have been used for millennia in Eastern traditional medicine as cancer remedies [(13,14)] and recent scientific publications have corroborated the presence of anticancer molecules in mushroom extracts. For instance, some polysaccharides from *Ganoderma* possess anticancer activity through immunomodulatory, anti-proliferative, pro-apoptotic, anti-metastatic, and anti-angiogenic effects [(15)]; lentinan, a [3-glucan from *Lentinus* is licensed as a drug for treating gastric cancer where it activates the complement system [(16)]; other [3-glucans from *Ganoderma* and *Grifola* [(17)] show also great potential as modulators of the immune system; lectins from *Clitocybe* provide distinct carbohydrate-binding specificities, showing immunostimulatory and adhesion-/phagocytosis-promoting properties [(18)]; also triterpenoids from *Inonotus* have been shown to induce apoptosis in lung cancer cell lines [(19)]. In addition, the immense number of mushroom species around the world [(20)], and their sessile nature, suggests that many fungal natural products may be still waiting for discovery.

Our goal in this study was to screen a number of scarcely known mushroom species collected in the rain forest of Costa Rica looking for new antitumor molecules. The most potent extract was found in specimens of *Macrocybe titans*, a local mushroom of very large proportions which usually grows in anthills of leaf cutter ants [(21)]. The molecule responsible for the anticancer activity was isolated, characterized, synthesized, and its anticancer properties were demonstrated both *in vitro* and *in vivo*.

## 2. Results

Several mushroom species were collected in the rain forest of Costa Rica, classified by expert botanists, and brought to the laboratory for extract preparation. Each specimen was subjected to several extraction procedures, including water- and ethanol-based techniques. All extracts were freeze-dried and sent to CIBIR for anticancer activity testing in cell lines. Several extracts from different species showed some “anticancer” properties, defined as the ability of destroying cancer cells while being innocuous to nontumoral cells (Fig. 1).

**Figure 1.**
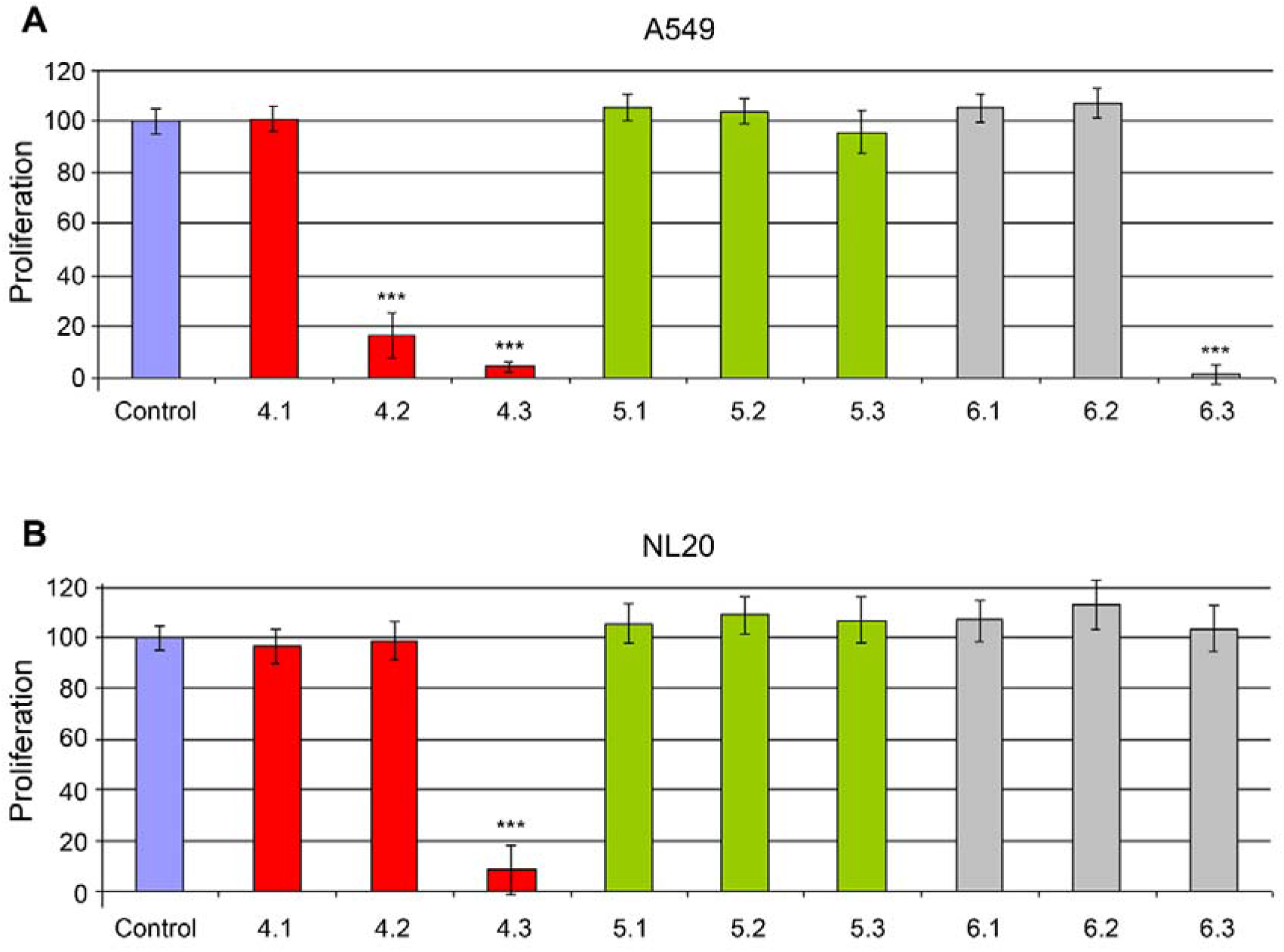
Toxicity of different mushroom extracts on cell lines A549 **(A)** and NL20 (B). All extracts were added at 0.3 mg/mL in growth medium containing 1% FBS, and incubated for 5 days. Comparing extract effects in both cell lines, three behaviors can be identified: i) no effect on either cell line (extract 4.1 for example), ii) toxic for both cell lines (extract 4.3), and iii) “antitumoral” effect (toxic for A549 and non-toxic for NL20, extracts 4.2 and 6.3). Bars represent the mean ± SD of 8 independent measures. ***: p<0.001 vs control.

The most promising sample was the precipitated phase of a 95% ethanol extract from the mushroom *Macrocybe titans*. Specifically, we began with 3.5 Kg of fresh mushroom specimens. These were dried to obtain 367.2 g of dry mushroom powder, which was extracted with 2 washes, 30 min each, in 500 mL 95% ethanol at room temperature, with ultrasounds. The ethanol extracts were kept at −20”C for 24 h and, then, the soluble phase was separated from the precipitate. This species was chosen for further analysis.

The initial extract was fractionated following different techniques and the resulting fractions were tested again in the cell lines to follow the “anticancer” activity. The finally successful strategy consisted in an initial alkaloidal separation followed by column chromatography with polymeric resin (HP20-SS) using different mobile phases. The best ratio of selectivity was obtained with an IPA:CH2Cl2 8:2 mobile phase at pH 12.0 (Fig. 2).

**Figure 2.**
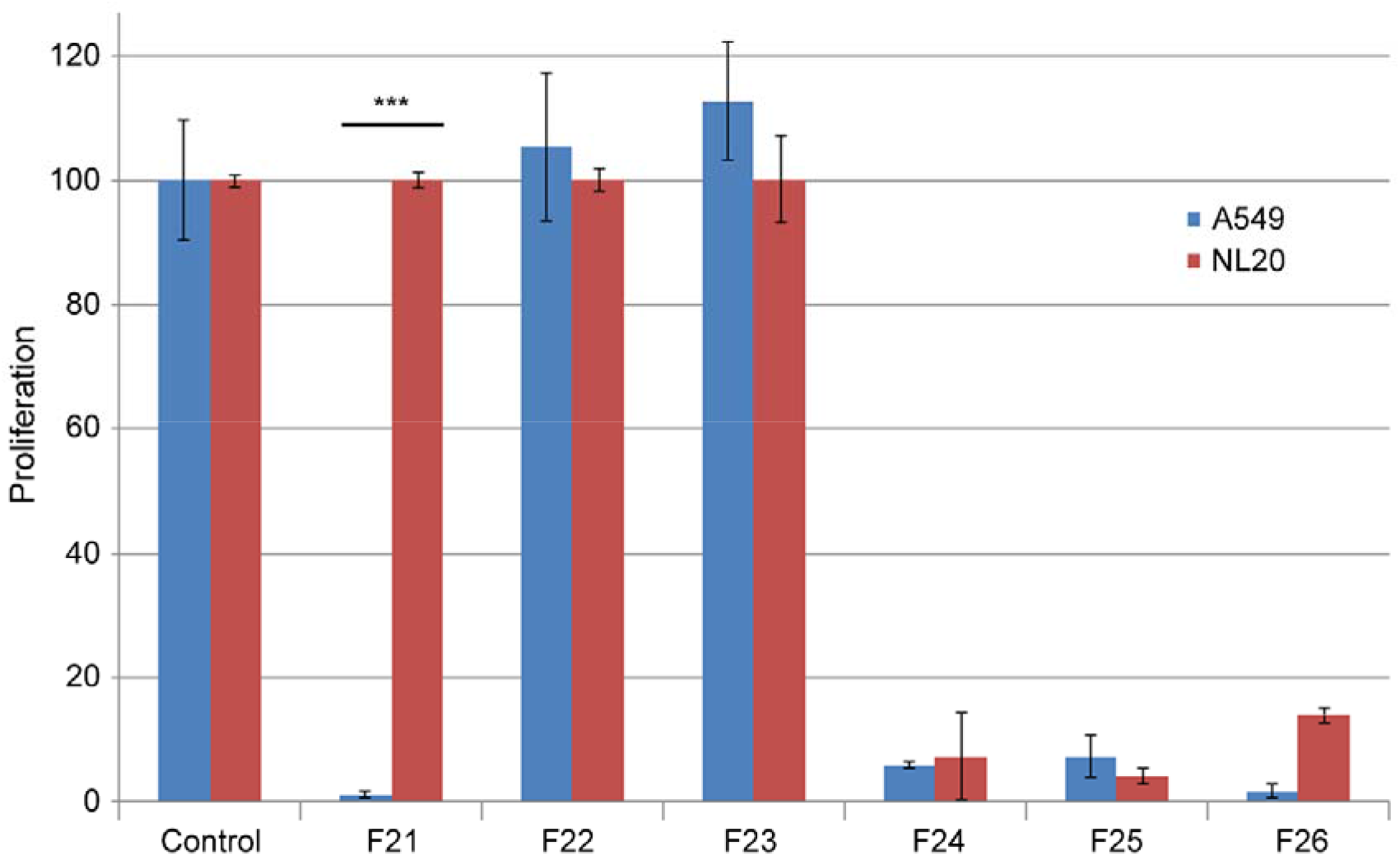
Toxicity of several fractions obtained by chromatography of the initial *M. titans* extract on cell lines A549 (blue bars) and NL20 (red bars). All extracts were added at 0.3 mg/mL in growth medium containing 1% FBS, and incubated for 5 days. Fraction 21 (F21) presents the desired characteristics of destroying the cancer cells while preserving the non-tumoral cells. Bars represent the mean ± SD of 8 independent measures. ***: p<0.001 between cell lines.

Once the molecule responsible for the differential growth inhibitory activity was pure enough, the final fraction was subjected to NMR, mass spectrometry, and 2D-COSY (Supplementary Materials, Natural extract characterization) to determine its chemical structure.

Data analysis identified our molecule as a triglyceride containing palmitic acid, oleic acid, and a more complex fatty acid of 18-20 carbons containing two double bonds. Since the final structure was impossible to determine, we decided to synthesize all compatible triglycerides (Table 1) and test them for biological activity with the cell lines. The presence of the two double bonds provides the molecule with optical activity. Some molecules were synthesized as racemic mixtures, but for others both enantiomers were synthesized and tested. From the 10 triglycerides that were synthesized (Table 1) only one exerted differential growth inhibitory activity, specifically the R enantiomer of TG (C18:2, 9z,12z; C16:0; C18:1, 9z) (Supplementary Materials, Growth modulatory activity of synthetic triglycerides). A search of the Chemical Abstract database came out negative, indicating this is a new structure. This triglyceride was named Macrocybin because of its origin from the *Macrocybe titans* mushroom.

**Table 1.** Triglycerides synthesized for this study and chemical structure of Macrocybin. The structure of the other triglycerides is shown in Supplementary Materials (Triglyceride synthesis).

| Internal code | Chemical formula |
| --- | --- |
| TG1 | TG (C16:0; C18:1, 9z; C20:2, 11z,14z) |
| TG2 | TG (C16:0; C20:2, 11z,14z; C18:1, 9z) |
| TG3 | TG (C16:0; C18:1, 9z; C18:1, 9z) |
| TG4 | TG (C18:0; C18:1, 9z; C18:2, 9z,12z) |
| TG5 | 2S-TG (C20:2, 11z,14z; C18:1, 9z; C16:0) |
| TG6 | 2R-TG (C20:2, 11z,14z; C18:1, 9z; C16:0) |
| TG7 | 2S-TG (C20:2, 11z,14z; C16:0; C18:1, 9z) |
| TG8 | 2R-TG (C20:2, 11z,14z; C16:0; C18:1, 9z) |
| TG9 | 2S-TG (C18:2, 9z,12z; C16:0; C18:1, 9z) |
| TG10 (Macrocybin) | 2R-TG (C18:2, 9z,12z; C16:0; C18:1, 9z) |

To determine the similarities between the synthetic molecule and the mushroom fraction, their NMR spectra were compared (Fig. 3) and they were very similar.

**Figure 3.**
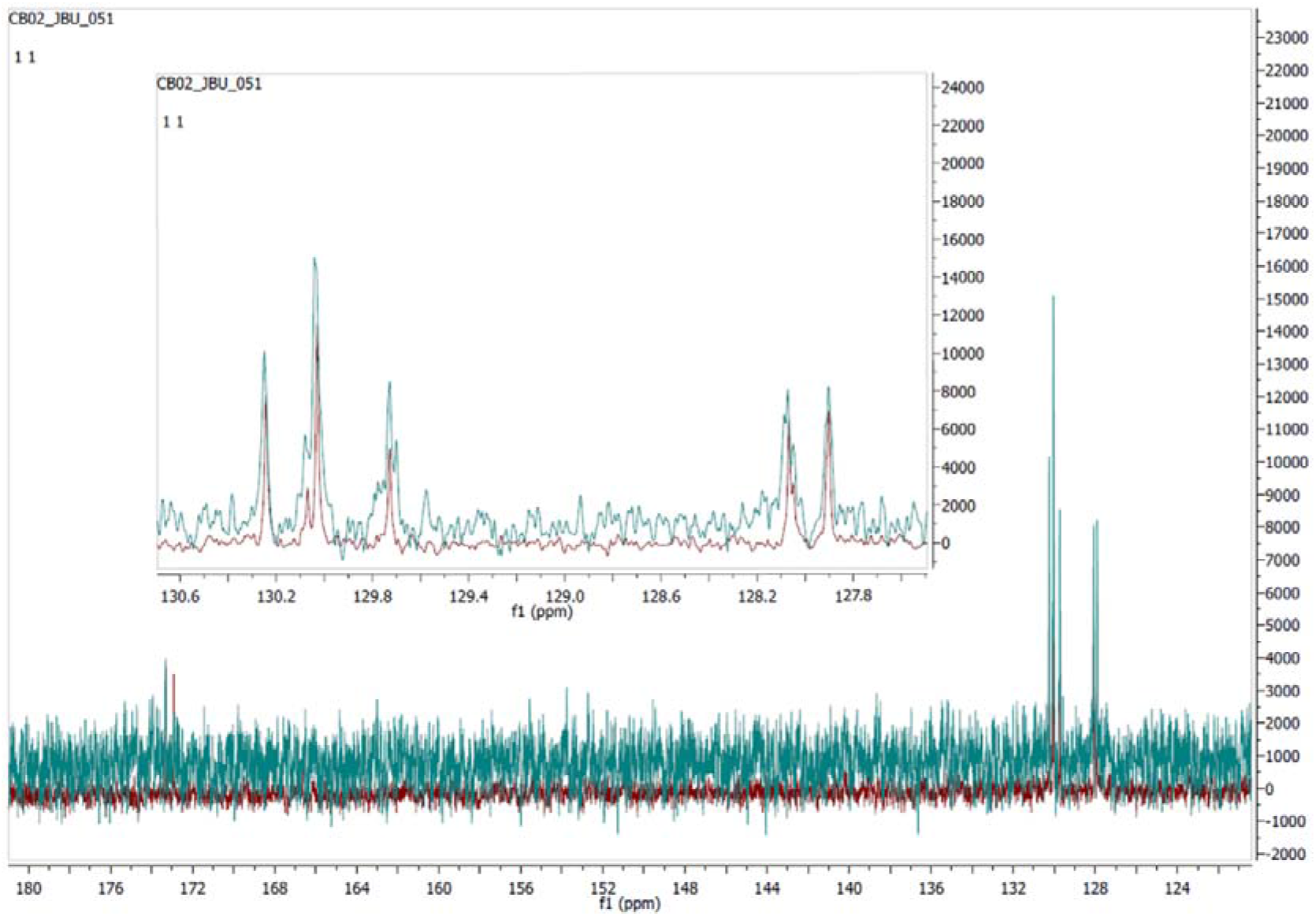
Comparison of ^13^C-NMR spectra of olefinic carbons of the synthetic triglyceride TG10 (red) and the natural extract (green). Spectra comparing the aliphatic carbons are shown in Supplementary Materials.

Toxicity assays for TG10 demonstrated a wide ratio of selectivity between A549 and NL20, with IC_50_ values of 13.4 and 50.1 μg/ml, respectively (Fig. 4), thus confirming the *in vitro* “anticancer” activity of the synthetic triglyceride, Macrocybin.

**Figure 4.**
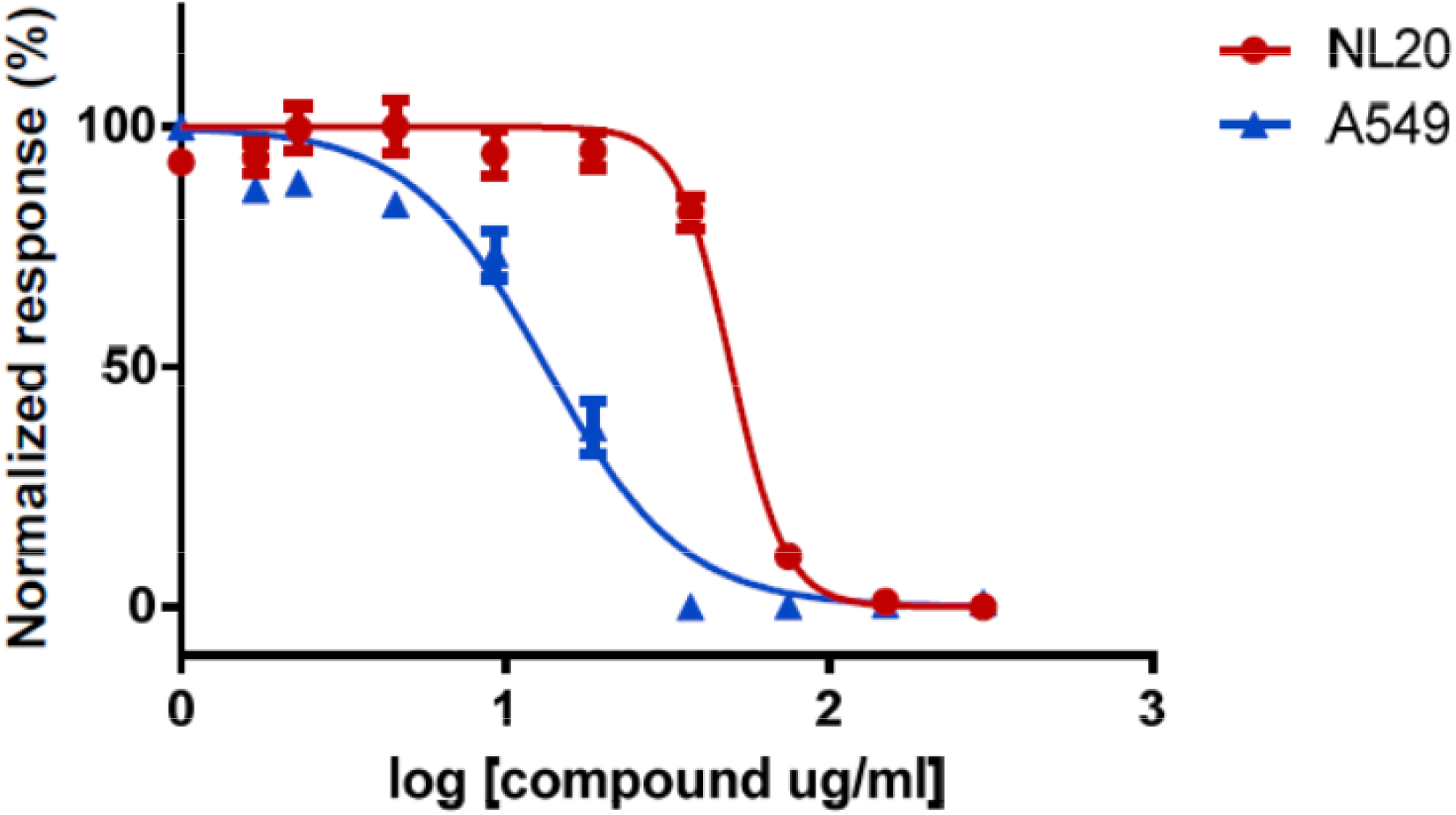
Concentration-dependent toxicity of synthetic Macrocybin on cell lines A549 (blue triangles) and NL20 (red circles). The synthetic try glyceride is more toxic for tumor cells than for non-tumoral cells. Error bars represent the SD of 8 independent measures.

To determine whether this new molecule had antitumor properties *in vivo*, a xenograft study was performed, using A549 as the tumor-initiating cell. Tumors of mice receiving vehicle injections grew progressively until they reached the maximum volume allowed for humane reasons and mice had to be sacrificed. On the other hand, tumors injected with Macrocybin grew somewhat more slowly (p<0.05 after 40 days of treatment), indicating a therapeutic function for the triglyceride (Fig. 5).

**Figure 5.**
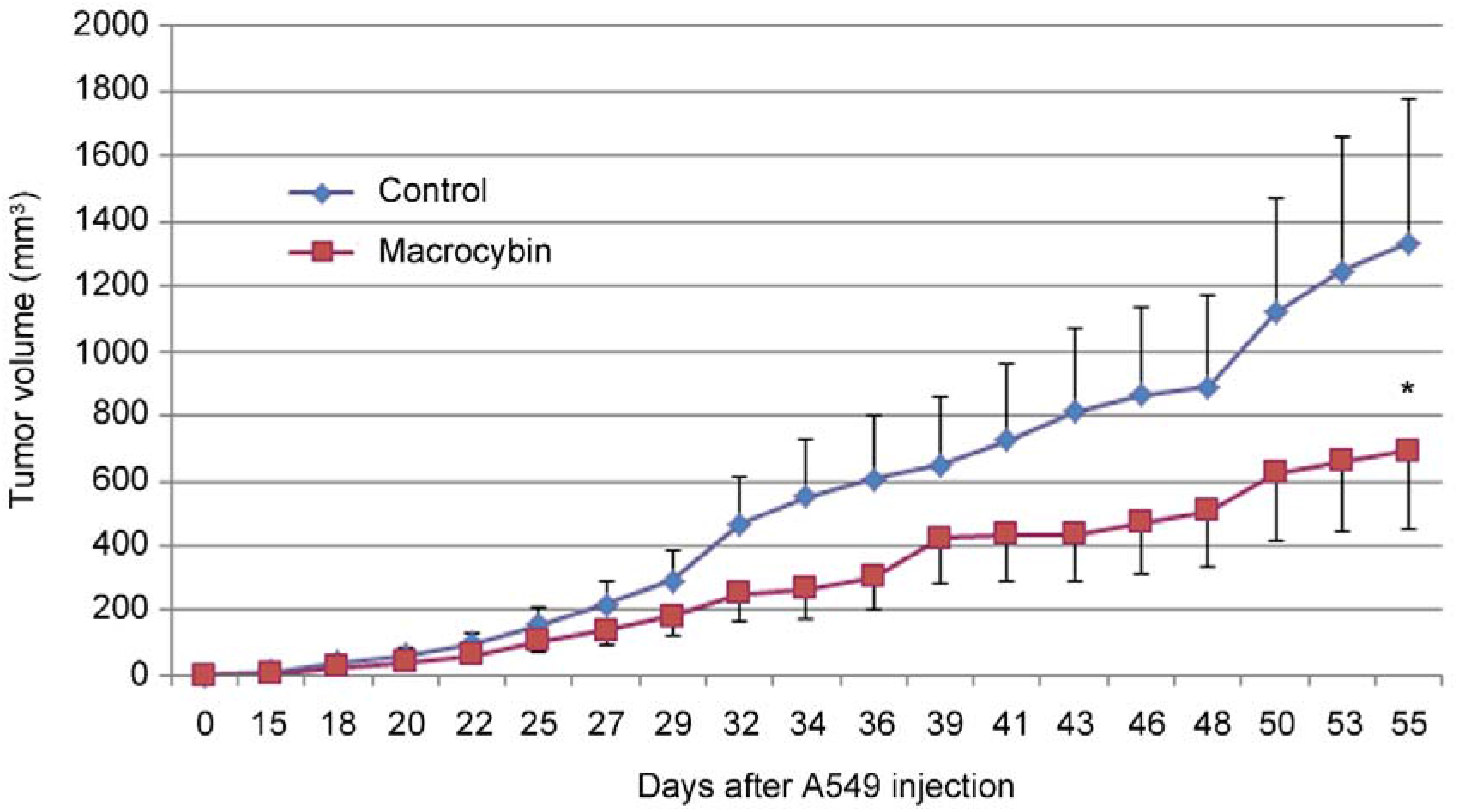
Xenograft experiment. Evolution of tumor volume in control (blue diamonds) and Macrocybin treated (red squares) A549 tumors grown in the flank of experimental mice. Cell injection was performed at day 0, and intratumoral injection began on day 15. There was a significant difference between treatments (ANOVA, p<0.05). Each point represents the mean ± SD for 10 mice. *: p<0.05 between treatments. Raw data values are shown in Supplementary Material.

To investigate the potential mechanism of action driving this antitumor activity, A549 and NL20 cells were stained with cytoskeleton-labeling moieties after exposure to Macrocybin. No differences were found in the tubulin cytoskeleton (results not shown) but the actin cytoskeleton, as stained with phallacidin, was dismantled in A549 tumor cells by the triglyceride whereas it was unaffected in NL20 cells. The actin molecules of A549 translocated to the cell membrane forming filopodia (Fig. 6).

**Figure 6.**
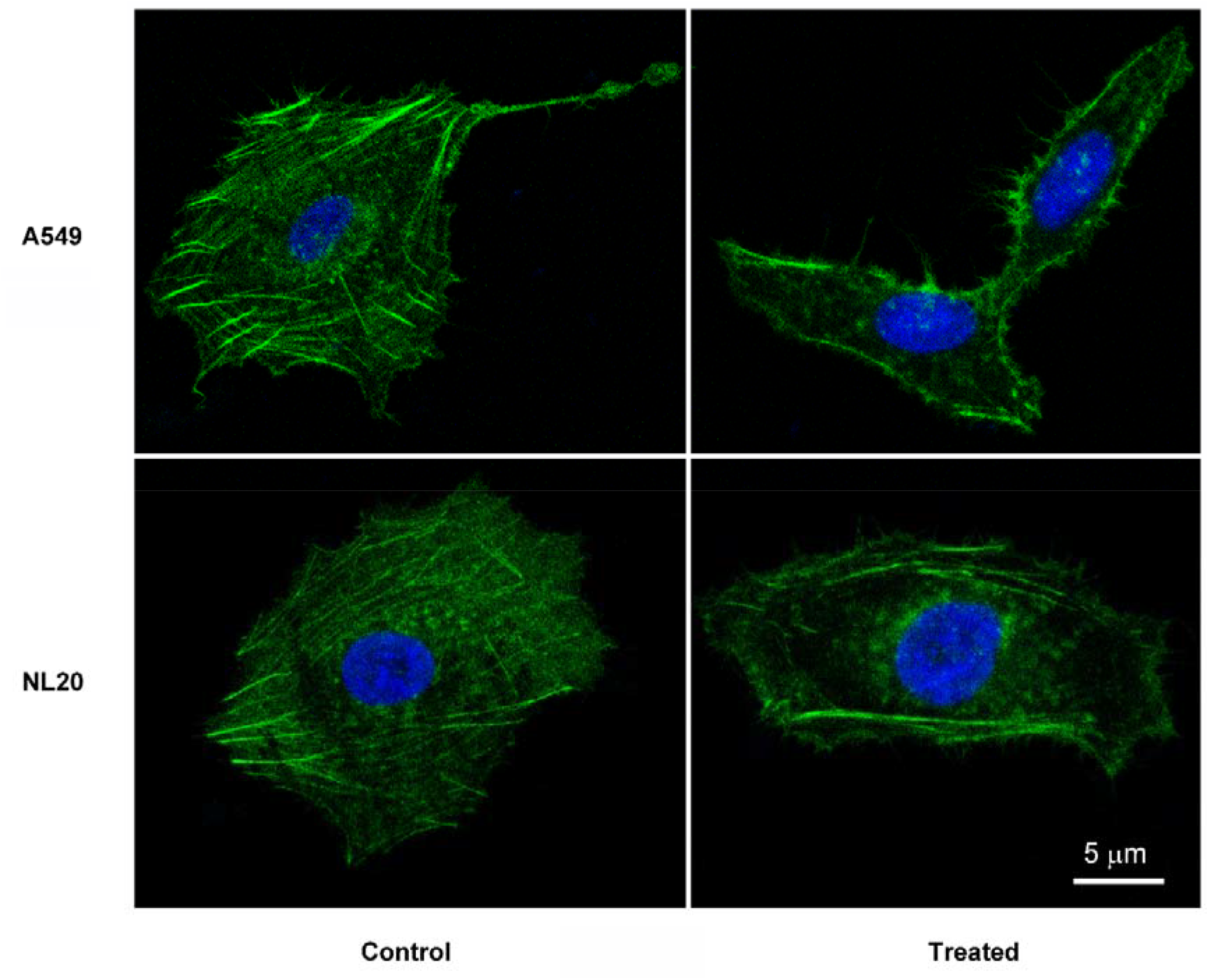
Representative confocal microscopy images of A549 and NL20 cells treated, or not (control), with 37 μg/ml Macrocybin for 24 h. The actin cytoskeleton was stained with Bodipyphallacidin (green) and the nuclei with DAPI (blue). Following treatment, A549 cells lose their stress fibers and actin migrates to the cell membrane to produce small filopodia. Scale bar = 5 μm.

Macrocybin is a complex lipid and, as such, could be internalized into the cell through a number of lipid transport mechanisms [(22)]. To investigate whether a preferential transport into tumoral cells takes place, we analyzed Macrocybin-induced changes in lipid transport molecules (Table 2) through qRT-PCR. A549 and NL20 cells were seeded in 6-well plates and Macrocybin (or PBS as vehicle) was added at 37 μg/ml for 6h. The only significant differences were found for Caveolin-1 whose expression increased significantly (p<0.05) in Macrocybin-treated A549 cells over the untreated controls. In contrast, Macrocybin did not affect Caveolin-1 expression in NL20 cells (Fig. 7).

**Figure 7.**
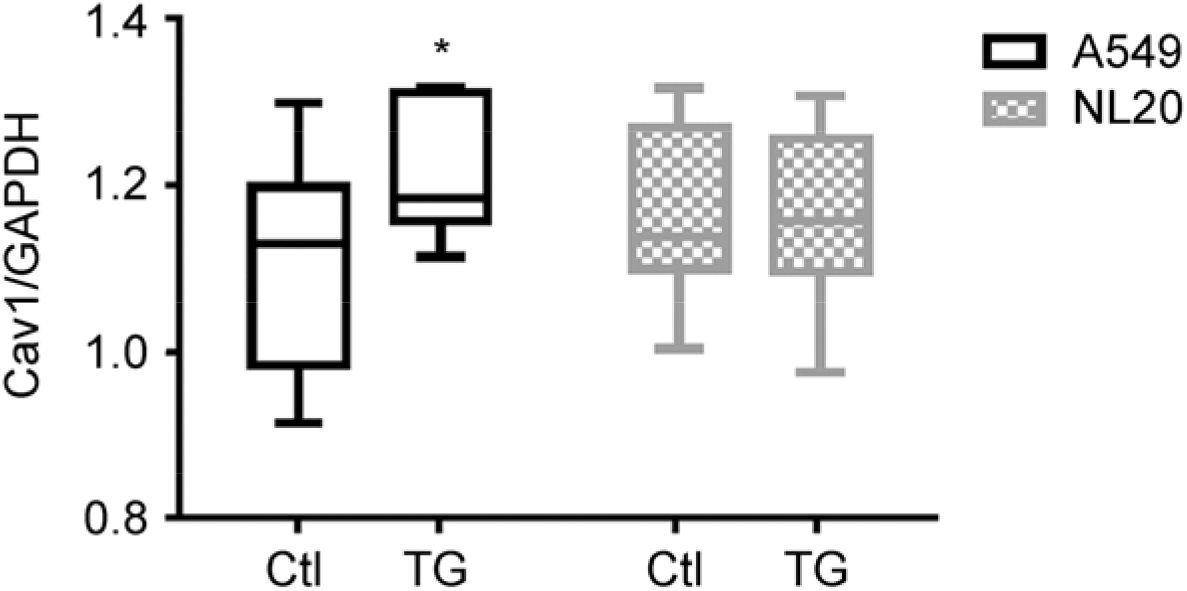
Gene expression for A549 and NL20 cells that were treated with 37 μg/ml Macrocybin (or control) for 6 h. Several lipid membrane transport proteins were analyzed by qRT-PCR. The only changes were observed on the expression of Caveolin-1 (Cav1) on treated A549 cells. Caveolin expression values were relativized to the housekeeping gene GAPDH. *: p<0.05 vs control (Ctl). This is a representative example of 3 independent experiments. Raw data values are provided in Supplementary Material.

**Table 2.**
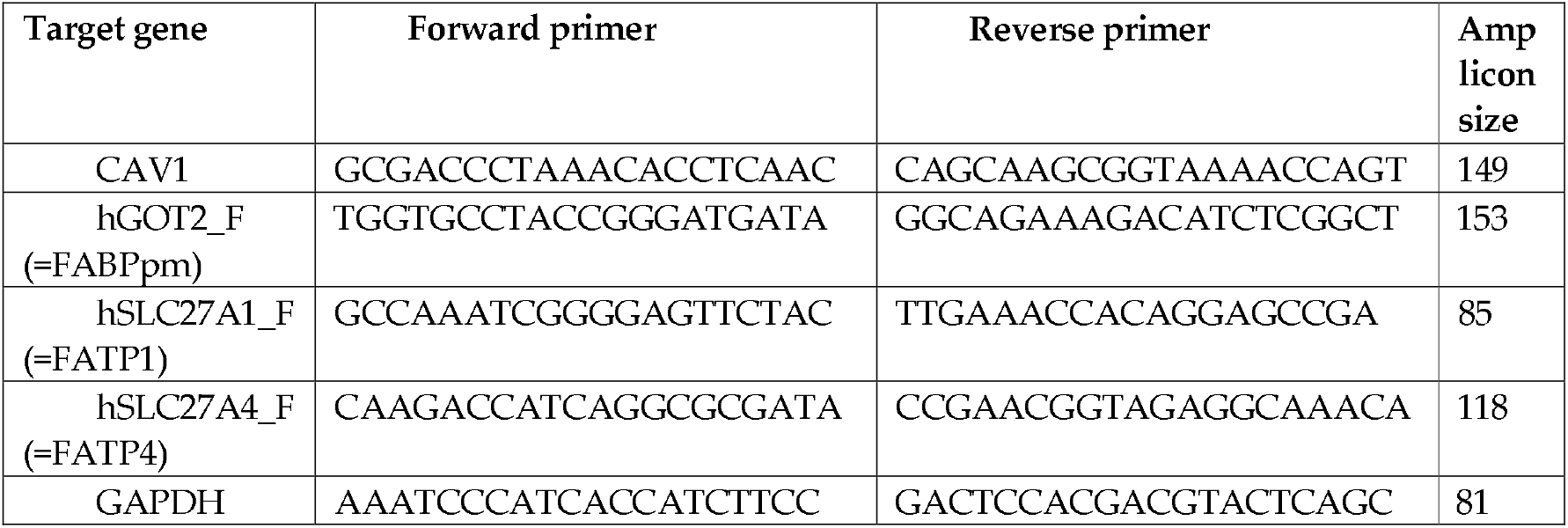
Primers used for qRT-PCR in this study. Annealing temperature was 60°C for all primers. GAPDH was used as a housekeeping gene.

## 3. Discussion

We have shown that the Costa Rican mushroom *Macrocybe titans* has anticancer properties. The molecule responsible for this activity was isolated to purity and identified as a complex triglyceride with formula 2R-TG (C18:2, 9z,12z; C16:0; C18:1, 9z). This molecule was named Macrocybin, was synthesized in the laboratory, and the synthetic molecule retained the anticancer properties both *in vitro* and in a xenograft model *in vivo*. The mechanism of action of the new molecule involves Caveolin-1 overexpression and actin cytoskeleton disorganization in the cancer cells.

The mushroom *M. titans* is a species which inhabits the tropical and subtropical regions of America and has been found between Florida and Argentina [(23)]. Other species of the same genus are found in tropical regions around the world, and include *M. gigantea* (India), *M. crassa* (Sri Lanka), *M. lobayensis* (Ghana), and *M. spectabilis* (Mauritius), among others [(21)]. *Macrocybe titans* produces arguably the largest carpophores in the world of fungi, with some specimens reaching 100 cm in diameter and weighting up to 20 Kg [(21)]. Interestingly, the specimens found in Costa Rica appear predominantly in the vicinity of the nests of gardening ants (*Atta cephalotes*) suggesting a potential symbiotic relationship between these two species [(21)]. All members of the *Macrocybe* genus are considered edible [(24)]. Although these mushrooms have not been reported as medicinal remedies in traditional pharmacopeia, recent studies have found that some polysaccharides of *M. titans* inhibit melanoma cell migration [(25)] and that the fruiting body of *M. gigantea* contains antimicrobial compounds [(26)]. These discoveries underscore the need for preserving biodiversity as a source for novel drugs and drug precursors [(27)].

The idea of a triglyceride acting as a therapy against cancer may seem rather counterintuitive. High levels of triglycerides in the blood constitute a clear risk for cancer initiation and progression since they provide a rich source of energy for developing tumors [(28)] and may increase metastatic potential [(29)]. Nevertheless, there are some specific types of cancer, such as breast [(30)] or prostate [(28)], where high levels of circulating triglycerides correlate with a better prognosis. In addition, a few studies have identified specific lipids with therapeutic functions. For instance, conjugated linoleic acid has been described as a natural anticarcinogenic compound [(31)].

Our studies testing the anticancer efficacy of different synthetic candidates indicate a high level of specificity in the anticancer actions of Macrocybin. Even the S enantiomer of Macrocybin was devoid of physiological activity. This indicates that the anticancer activity is dependent on a very specific interaction of Macrocybin with biological components (receptors and/or membrane transporters) of the tumor cells, rather than a bulk role such as energy provider (which generic blood triglycerides may play).

Macrocybin was able to reduce tumor growth in a xenograft model of human lung cancer. As a triglyceride, Macrocybin has a modest solubility in water-based vehicles. Perhaps future formulations using micelles [(32)] or nanoparticles [(33)] may increase Macrocybin biodisponibility and antitumor efficacy. Also, combinations of Macrocybin with other chemotherapeutic or immunotherapeutic drugs may represent promising avenues to reduce tumor burden [(34)].

Our results show that Macrocybin affects differentially the actin cytoseleton of tumor vs nontumor cells, and that this mechanism is mediated by changes on the expression of Caveolin-1. Caveolin-1 has been shown as a lipid transporter in a variety of cell types [(35–37)]. Changes in the expression of this protein have been related with lung cancer prognosis [(38)], although some studies suggest that this relationship may be context-dependent [(39)]. This protein may constitute the entry point for Macrocybin into the cell and the fact that its expression is affected in tumor cells but not in nontumor cells may partially explain the preferential toxicity of the triglyceride for cancer cells.

Disruption of the actin cytoskeleton is a common mechanism of action for many natural products with antitumor properties. For instance, proteoglucans extracted from *Grifola* mushrooms affect actin cytoskeleton rearrangements, thus decreasing breast cancer cell motility [(40)]. Cucurbitacin I disrupts actin filaments in A549 cells, resulting in a reduction of lung cancer cell growth [(41)]. Enterolactone, a flaxseed-derived lignan, alters FAK-Src signaling and disorganizes the actin cytoskeleton, suppressing migration and invasion of lung cancer cells [(42)]. Another example is narciclasine, an isocarbostyril alkaloid isolated from *Amaryllidaceae* plants, which impairs actin cytoskeleton organization and induces apoptosis in brain cancers [(43)]. All these studies suggest that interfering with the actin cytoskeleton of tumor cells provides a useful approach to induce tumor cell death and to prevent metastasis by reducing tumor cell motility.

Given the ubiquity among cancer cells of Caveolin-1 and the actin cytoskeleton, Macrocybin may constitute a common therapeutic agent for a variety of cancers. Many further investigations will be necessary to evaluate in more detail the usefulness of Macrocybin as an antitumoral drug or lead structure. These studies must include, among others, studies on the systemic applicability. Future studies would also have to investigate whether other tumors are also susceptible to this new antitumor compound.

## 4. Materials and Methods

### 4.1. Mushroom collection and extract preparation

Several mushroom specimens belonging to different species were collected in the rain forests of Costa Rica by expert personnel of Institute Nacional de Biodiversidad de Costa Rica (INBio), under a specific permit issued by the Costa Rican Government (Oficina Técnica de la Comisión Nacional de la Gestión de la Biodiversidad, CONAGEBIO). Mushroom fragments were subjected to different extraction procedures, including crude extracts, extraction in hot water (80°C) for several 30 min incubations, or in 95% ethanol at 50°C for several 30 min incubations, with different exposures to ultrasound treatments. Final fractions, including the pellet and supernatant of each extraction, were freeze-dried and sent to CIBIR for screening. Voucher samples of the fungal collection are kept at INBio (Santo Domingo, Heredia, Costa Rica).

### 4.2. Anticancer screening strategy (toxicity assays)

Two human cell lines were used to screen the anticancer properties of the mushroom extracts: lung adenocarcinoma A549, and non-tumoral immortalized bronchiolar epithelial cell line NL20 (ATCC). Both cell lines were exposed in parallel to different concentrations of particular extracts for 5 days, and cytotoxicity was measured by the MTS method (Cell Titer, Promega), as reported [(44)]. To identify a sample as having “anticancer° properties it had to destroy cancer cells while preserving non-tumoral cells (or at least showing a wide therapeutic window between both lines). This strategy was used iteratively to guide extract purification until the final molecule could be identified.

Following ATCC’s protocols, A549 cells were maintained in Ham’s F12 medium containing 10% fetal bovine serum (FBS) and NL20 cells in complete medium (Ham’s F12 with 2 mM L-glutamine, 1.5 g/L sodium bicarbonate, 2.7 g/L D-glucose, 0.1 mM NEAA, 0.005 mg/ml insulin, 10 ng/ml EGF, 0.001 mg/ml transferrin, 500 ng/ml hydrocortisone, and 4% FBS).

Before starting the project, optimization of the toxicity assay was performed, testing different numbers of cells per well, days of incubation with the extracts, and FBS contents. We wanted all cells receiving exactly the same amount of extracts in the same medium, so we chose the NL20 medium which is the most complex and restrictive. FBS concentration was reduced to 1% to allow for a 5 day incubation period. Specifically, A549 and NL20 were seeded in 96-well plates at different densities (2,000 and 10,000 cells/well, respectively) in complete NL20 medium containing 1% FBS, in a final volume of 50 μl/well, and incubated at 37°C in a humidified atmosphere, containing 5% CO2. The next day, extracts were added at the indicated concentrations, in other 50 μl/well, in the same medium, and incubated for 5 days. At the end of this period, 15 μl of Cell Titer were added per well and, after an additional incubation of 4 h, color intensity was assessed in a plate reader (POLARstar Omega) at 490 nm.

To determine IC_50_ values, treatments were diluted at 0.6 mg/mL in test medium and 8 serial double dilutions were prepared. These solutions were added to cells and incubated as above. For each concentration point, 8 independent repeats were performed. Graphs and IC_50_ values were obtained with GraphPad Prism 8.3.0 software.

Cell lines were authenticated by STR profiling (IDEXX BioAnalytics).

### 4.3. Fractionation strategies

The mushroom with most anticancer activity was chosen for further analysis. The initial extract was subjected to different fractionation protocols, which included different chromatographic techniques. Every time that new fractions were separated, they were analyzed for their anticancer properties through cell line screening (as above) and the positive ones were subjected to further fractionation, until obtaining a pure enough compound that could be identified through analytical chemistry methods.

Many combinations were tested, but the one that resulted in final successful purification included alkaloidal separation followed by column chromatography with polymeric resin HP20-SS, using different mobile phases.

### 4.4. Elucidation of the compound’s chemical structure

The final extract was subjected to analytical techniques including ^1^H-nuclear magnetic resonance (NMR), ^13^C-NMR, DEPT-135-NMR, mass spectroscopy (Q-TOF), and bidimensional experiments HMQC and COSY. Interpretation of the data indicated that the new compound was a triglyceride and the component fatty acids were identified to a certain degree of confidence, but the exact location of the fatty acids in the triglyceride and the position of the two double bonds in the more complex fatty acid could not be completely determined. Therefore we synthesized the most probable molecules (Table 1) and tested them with our cell line screening strategy. Given the presence of two double bonds in one of the fatty acids, the triglyceride possesses optical activity, so both enantiomers were synthesized (Table 1) and tested.

### 4.5. Triglyceride synthesis

Molecules in Table 1 were synthesized following standard techniques. Details are shown in Supplementary Materials (Triglyceride synthesis).

### 4.6. Anticancer activity in a xenograft model

Once the active synthetic molecule (Macrocybin) was available, it was tested *in vivo* following standard protocols [(45)]. Briefly, 10×10^6^ A549 cells were injected in the flank of twenty 8-week old NOD *scid* gamma mice (NSG, Stock No. 005557, The Jackson Laboratory). Animals were randomly divided in 2 experimental groups and, when tumors became palpable, they were injected intratumorally with 100 μl of vehicle (PBS) in the control group (n=lO), or with 0.1 mg/ml Macrocybin in 100 μl of PBS in the treatment group (n=lO), three times a week. Just before injection, tumor volume was estimated with a caliper by measuring the maximum length and width of the tumor and applying the formula: Volume = (width)^2^ x length/2 [(46)]. When tumor volume reached 2,000 mm^3^, mice were sacrificed for ethical considerations.

All procedures involving animals were carried out in accordance with the European Communities Council Directive (2010/63/UE) and Spanish legislation (RD53/2013) on animal experiments and with approval from CIBIR’s committee on ethical use of animals (Órgano Encargado del Bienestar Animal del Centro de Investigación Biomédica de La Rioja, OEBA-CIBIR).

### 4.7. Cytoskeleton staining and confocal microscopy

To test whether Macrocybin, like other natural products, acts through cytoskeleton interactions, A549 and NL20 cells were seeded in 8-well chamber slides (Lab-Tek II), treated with different concentrations of Macrocybin (or vehicle) for 24 h, washed, fixed with 10% formalin, and permeabilized for 10 min with 0.1% Triton X-100. For cytoskeleton imaging, cells were exposed to 1:1,000 mouse antitubulin antibody (T6O74, Sigma-Aldrich) overnight at 4°C, washed with PBS, incubated with a mixture of 1:200 Bodipy-phallacidin (Molecular Probes) and 1:400 goat-anti mouse IgG labeled with Alexa Fluor 633 (A-21052, Invitrogen), and washed again. Nuclear staining was achieved with DAPI (ProLong Gold Antifade Mountant, Invitrogen). Slides were observed with a confocal microscope (TCS SP5, Leica).

### 4.8. Gene expression

To identify cellular pathways potentially involved in Macrocybin action, mRNA was extracted from treated and untreated A549 and NL20 cells using the RNeasy MiniKit (Qiagen). Total RNA (1 μg) was reverse transcribed using the SuperScriptR III reverse Transcriptase kit (Thermo Fisher Scientific), and quantitative real-time PCR was performed as described [(47)]. Specific primers are shown in Table 2. GAPDH was used as a housekeeping gene.

### 4.9. Statistical analysis

All datasets were tested for normalcy and homoscedasticity. Normally distributed data were evaluated by Student’s t test or by ANOVA followed by the Dunnet’s post-hoc test while data not following a normal distribution were analyzed with the Kruskal-Wallis test followed by the Mann-Whitney U test. All data were analyzed with GraphPad Prism 8.3.0 software and were considered statistically significant when p<0.05.

## 5. Conclusions

In conclusion, we have shown that the mushroom natural product, Macrocybin, is a new biologically active compound which reduces tumor growth by disassembling the actin cytoskeleton, providing a potential new strategy to fight cancer.

## Supplementary Materials

The following are available online at www.mdpi.com/xxx/s1, Natural extract characterization, triglyceride synthesis, growth modulatory activity of synthetic triglycerides, spectral comparison of aliphatic components, and actual values for xenograft and qRT-PCR studies.

## Author Contributions

Conceptualization, M.V. and A.M.; methodology, J.G.-S., L.O.-C., A.L.-R., and J.B.-U.; formal analysis, J.G.-S., L.O.-C., A.L.-R., and J.B.-U; investigation, M.V., J.G.-S., L.O.-C., A.L.-R., and J.B.-U.; data curation, J.G.-S., L.O.-C., J.B.-U., and A.M.; writing—original draft preparation, A.M.; writing—review and editing, A.M.; supervision, A.M. All authors have read and agreed to the published version of the manuscript.

## Funding

This research was funded by Fundación Rioja Salud (FRS), la Agencia de Desarrollo Económico de La Rioja (ADER), project 20’17-I-IDD-00067, and Fondos FEDER.

## Acknowledgments

We gratefully acknowledge the work of Dr. Kattia Rosales and her colleagues collecting specimens at the Instituto Nadonal de Biodiversidad de Costa Rica (INBio).

## Conflicts of Interest

The authors declare no conflict of interest. The funders had no role in the design of the study; in the collection, analyses, or interpretation of data; in the writing of the manuscript, or in the decision to publish the results.

## Sample Availability

Samples of Macrocybin are available from the authors, while supplies last.

## Notes

### Competing Interest Statement

The authors have declared no competing interest.

