## Supplementary Material for "Macrocybin, a mushroom natural triglyceride, reduces tumor growth *in vitro* and *in vivo* through caveolin-mediated interference with the actin cytoskeleton"

Index

| | Page |
|---|---|
| 1. Natural extract characterization | 2 |
| 2. Triglyceride synthesis | 5 |
| 3. Growth modulation activity of synthetic triglycerides | 22 |
| 4. Comparison of aliphatic carbons between the synthetic triglyceride and the natural extract | 25 |
| 5. Dataset for xenograft study | 26 |
| 6. Datasets for qRT-PCR study | 29 |

### 1. Natural extract characterization

[1]H-NMR:

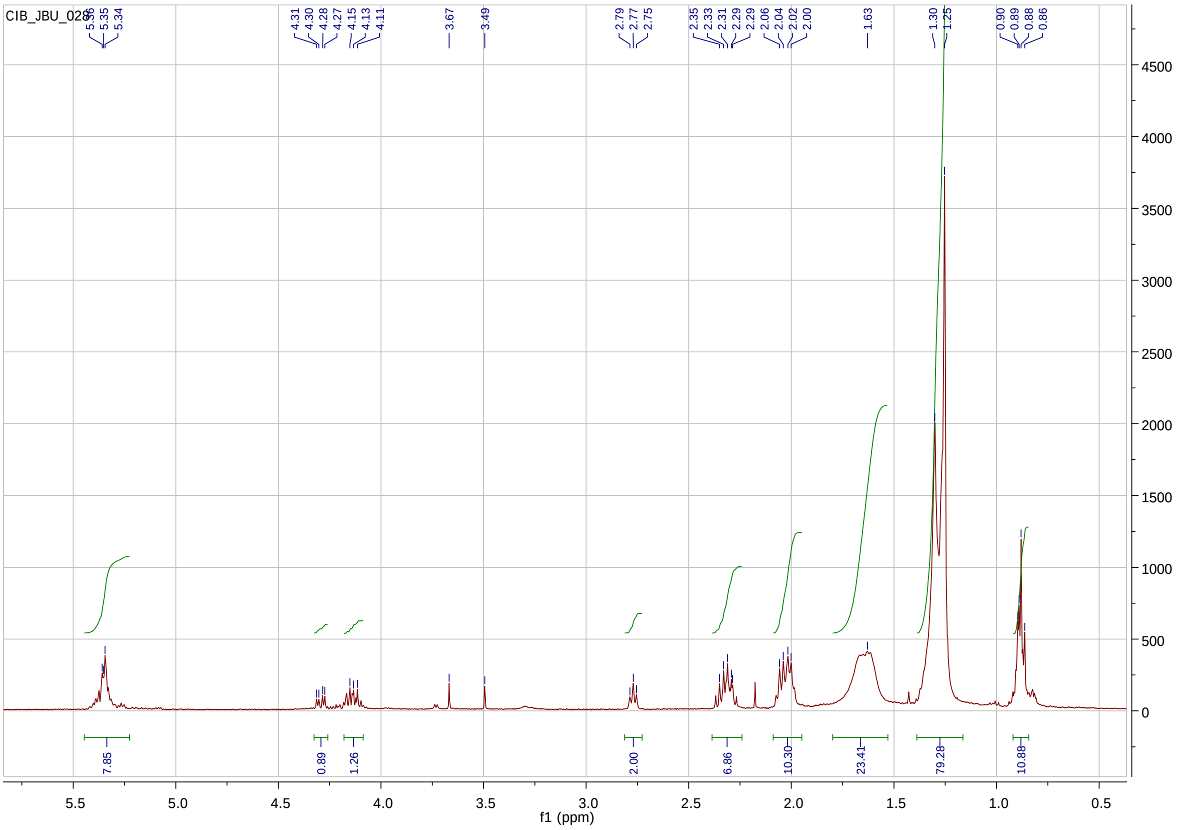

$^{13}$C-NMR:

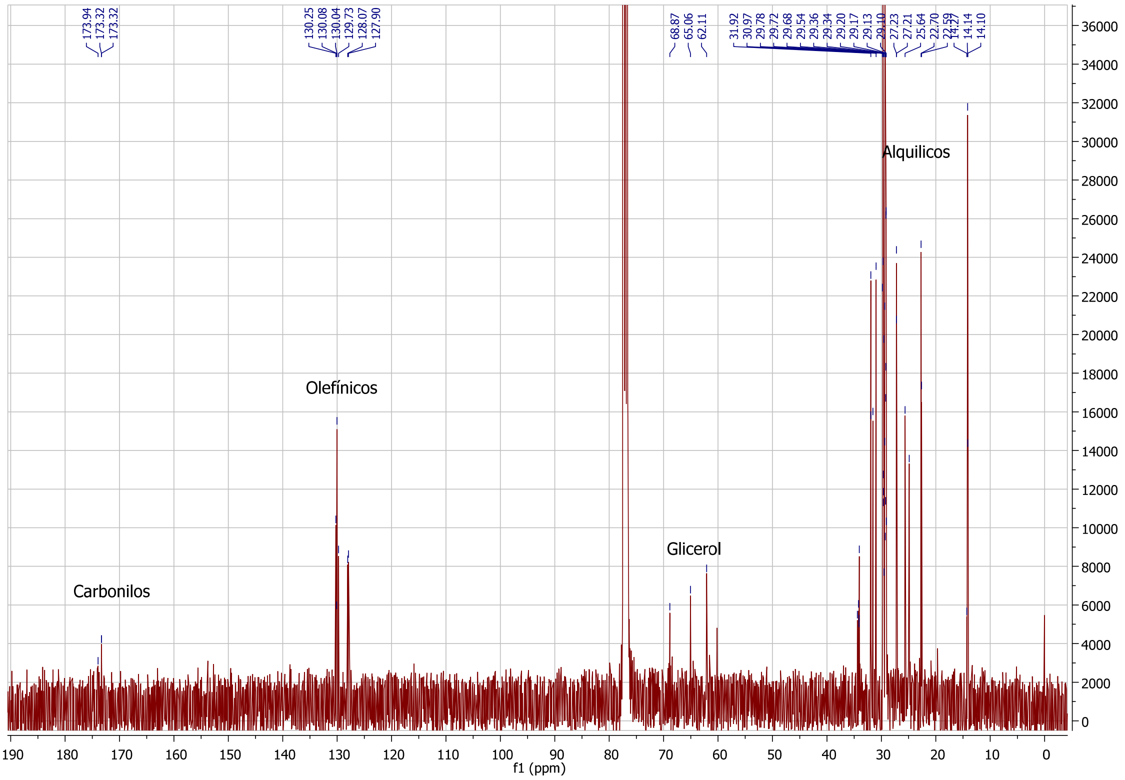

DEPT 135

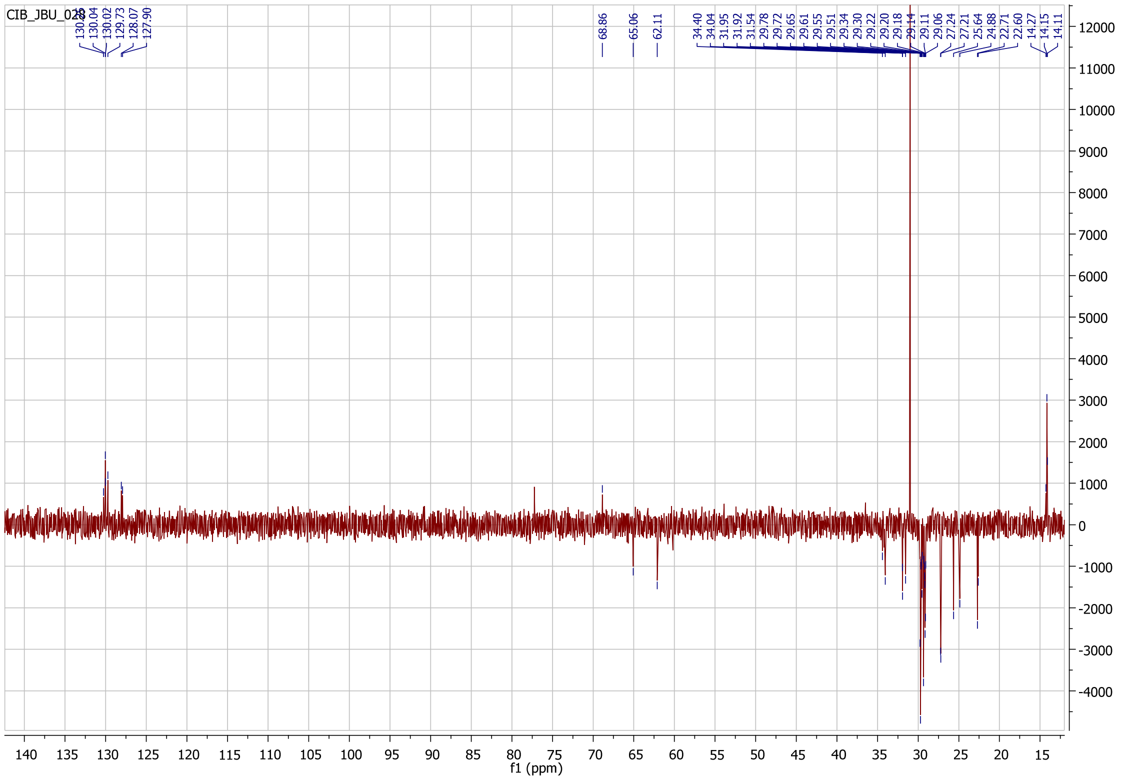

2D-COSY

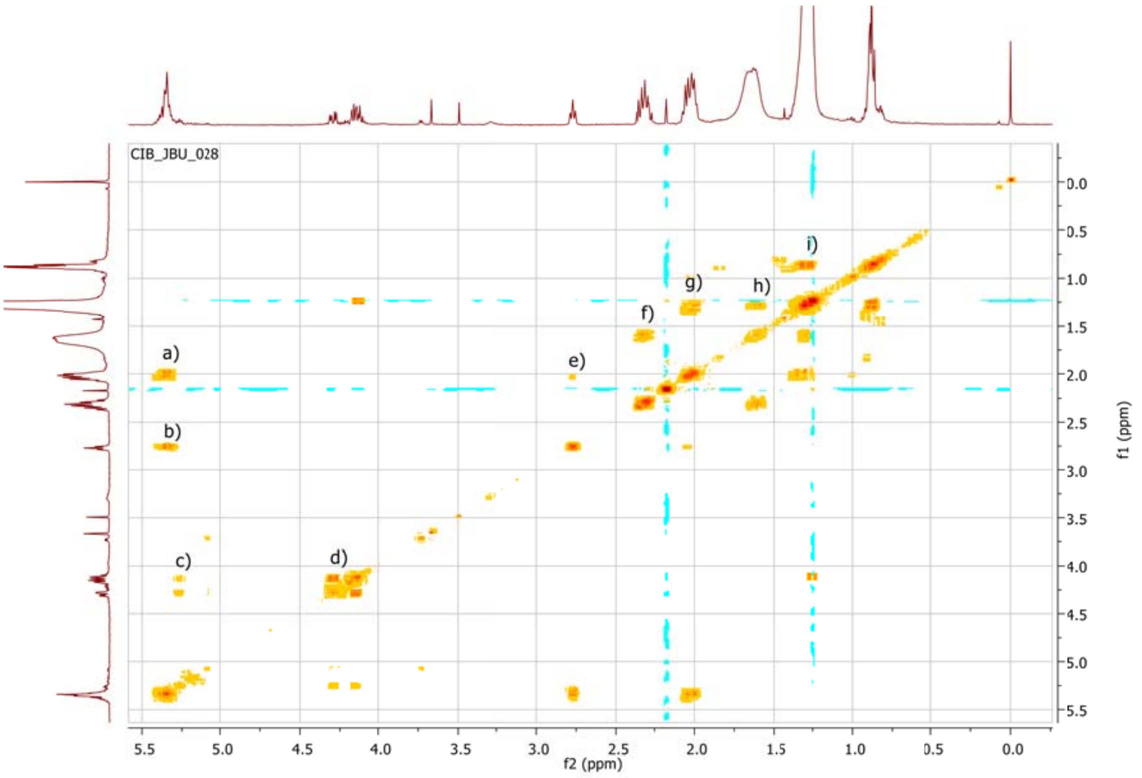
CIB_JBU_028
a)
b)
c)
d)
e)
f)
g)
h)
i)
5.5
5.0
4.5
4.0
3.5
3.0
2.5
2.0
1.5
1.0
0.5
0.0
f2 (ppm)
f1 (ppm)

### 2. Triglyceride synthesis

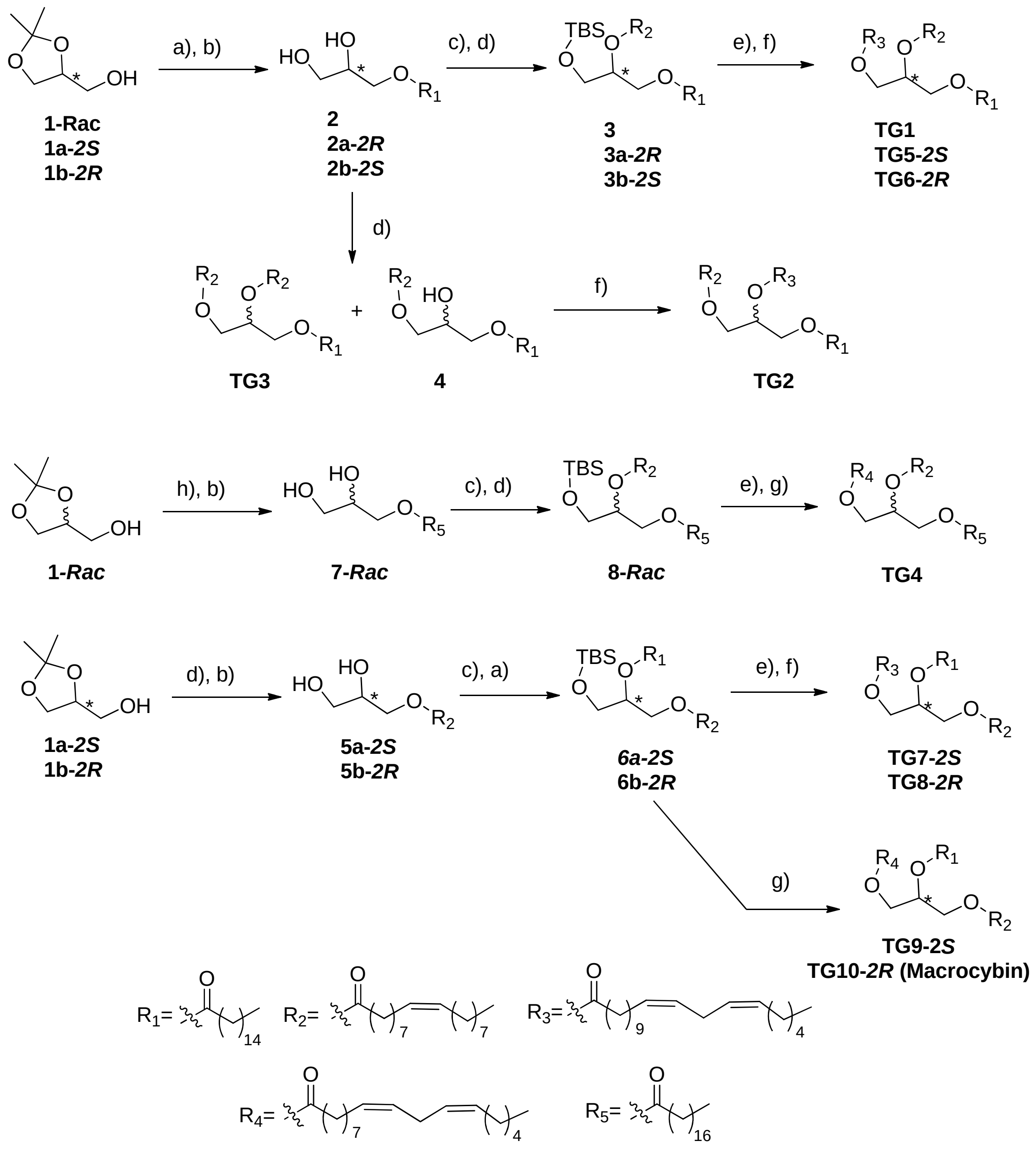

a) Palmitoyl chloride, py, DCM, 0ºC to rt, b)AcOH/$H_2O$ 60ºC, c)TBSCl, Imid, THF, 0ºC to rt, d) oleyl chloride, py, DCM, 0ºC to rt, e) HF·py, py, THF, 0ºC to rt, f) (*11Z,14Z*)-icosa-11,14-dienoic acid, EDCI, DMAP, $CHCl_3$, rt, g) linoleyl chloride, py, DCM, 0ºC to rt, h) stearyl chloride, py, DCM, 0ºC to rt.

**Experimental procedure A**. Ester formation from acylchloride.

To a solution of the starting alcohol in DCM (8 vol) at 0ºC, pyridine (5 eq) is added followed by the acyl chloride (1.3 eq). The mixture is left at rt overnight. 0.1 N HCl is added (8 vol) and stirred for 5 min at room temperature. The aqueous layer is discarded, and the organic layer is washed with water and brine, dried over anhydrous MgSO4 filtered and concentrated in vacuo. The crude is used in the following step without further purification.

**Experimental procedure B.** Acetal hydrolysis.The crude obtained in the step A is suspended in Acetic acid (10 vol) and water (2 vol) and stirred at 60ºC for 3 h. The acetic acid is evaporated under reduced pressure and the aqueous layer is extracted wiDCM. The organic phase is washed with water, 5% sodium bicarbonate in water, more water and brine. The DCM is dried over anhydrous MgSO4 filtered and concentrated in vacuo. The crude of the monoglyceride is used without further purification.

**Experimental procedure C**. Alcohol protection with Tert-Butyldimethylsilane.

To a solution of the monoglyceride obtained with the procedure B in THF (10 vol) at 0ºC imidazole (1.5 eq) is added followed by Tert-butyldimethylsilyl chloride (1.25eq). The mixture is stirred at rt overnight. The solid is filtered and washed with more THF and the solvent is removed under reduced pressure. The crude is used without further purification.

**Experimental procedure D**. Alcohol deprotection, silyl cleavage.

To a solution of the diglyceride silylprotected in THF (40 vol) at 0ºCpyridine (3 eq) is added followed by hydrogen fluoride pyridine (3 eq). The reaction is allowed to reach rt. After 3h at rt the mixture is poured into a 5% aqueous solution of sodium bicarbonate. The THF is removed under reduced pressure and the aqueous phase is extracted with MTBE. The organic layer is washed with 1N HCl, water and brine, dried over anhydrous MgSO4 filtered and dried in vacuo. The crude is used without further purification.

**Experimental procedure E.** Ester formation from the acid and alcohol.

To a solution of the alcohol and the corresponding acid (1.5 eq) in DCM at rt DMAP (30 eq) is added followed by the EDCI (20 eq). The mixture is stirred at rt for 16h. The solid is filtered and washed with more DCM. The solvent is removed under reduced pressure and the crude is purified by flash chromatography using hexane/MTBE mixture as eluent.

**TG (C16:0; C18:1, 9z; C20:2, 11z,14z),** ***(11Z,14Z)-2-(oleoyloxy)-3-(palmitoyloxy)propyl icosa-11,14-dienoate,*** **TG1.**

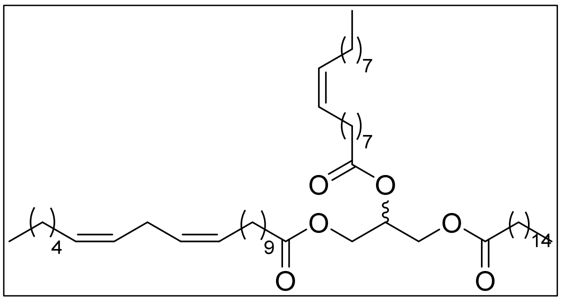

From 200 mg of (±)-(2,2-dimethyl-1,3-dioxolan-4-yl)methanol, following the **procedure A** with palmitoyl chloride, and **procedure B** to obtain **2** with quantitative yield. **2** is transformed following the

**procedure C**, followed by **procedure A** with oleoyl chloride to obtain 1.08 g of **3** in quantitative yield (2 steps). 95 mg of **TG1** are obtained from 112 mg of **3** following the **procedure D** and **procedure E** with (*11Z,14Z*)-icosa-11,14-dienoic acid after purification by flash chromatography using Hexane/MTBE 3% as eluent (72% yield, 2 steps). $^{1}$H NMR (400 MHz, $CDCl_3$) δ 5.42 – 5.28 (m, 6H), 5.26 (td, *J* = 5.9, 3.0 Hz, 1H), 4.29 (dd, *J* = 11.9, 4.3 Hz, 2H), 4.14 (dd, *J* = 11.9, 5.9 Hz, 2H), 2.77 (t, *J* = 6.4 Hz, 2H), 2.31 (td, *J* = 7.6, 2.4 Hz, 6H), 2.10 – 1.94 (m, 8H), 1.67 –1.57 (m, 6H), 1.41 – 1.18 (m, 62H), 0.93 – 0.84 (m, 9H). $^{13}$C NMR (101 MHz, $CDCl_3$) δ 173.4, 173.4, 173.0, 130.3, 130.3, 130.2, 129.9, 128.1, 128.1, 69.0, 62.2, 32.1, 32.0, 31.7, 30.3, 29.9, 29.8, 29.8, 29.8, 29.8, 29.7, 29.7, 29.6, 29.6, 29.5, 29.5, 29.4, 29.4, 29.3, 29.2, 27.4, 27.4, 27.3, 27.3, 25.8, 25.0, 25.0, 22.8, 22.7, 14.3, 14.2. Anal. Calcd for C57H104O6 (885.45 g/mol): C, 77.32; H, 11.84%. Found: C, 77.37; H, 11.90%.

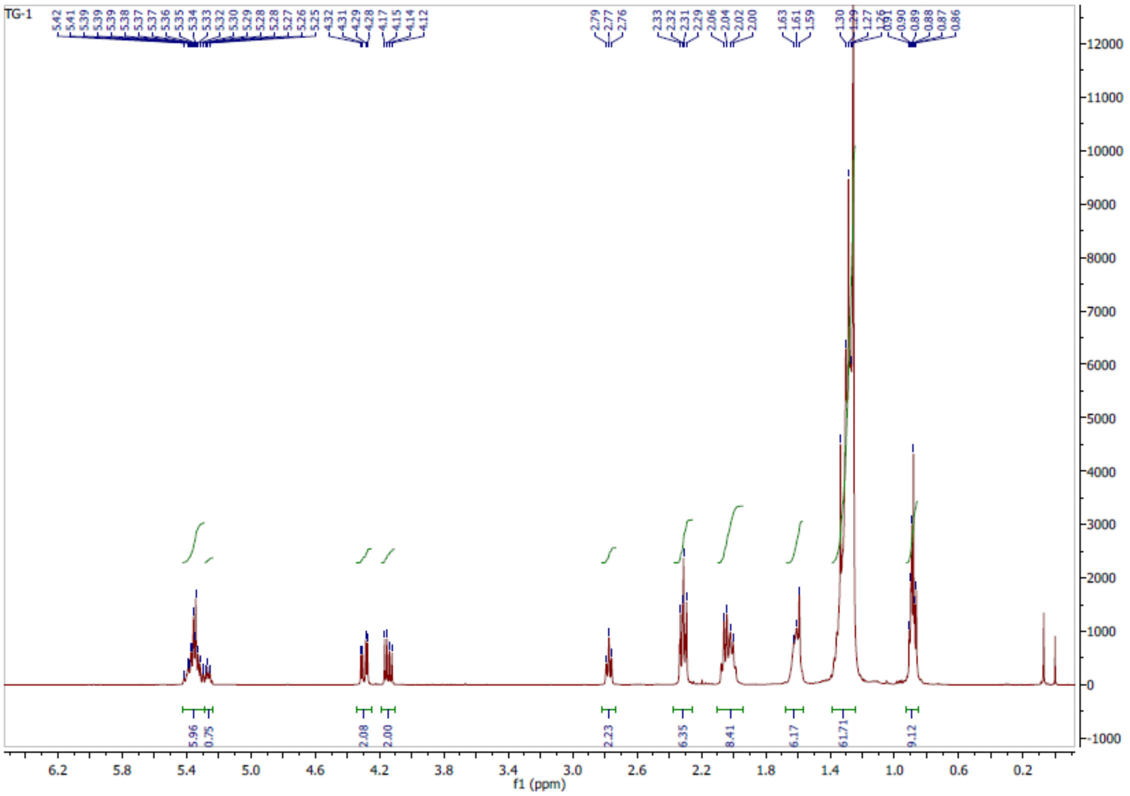

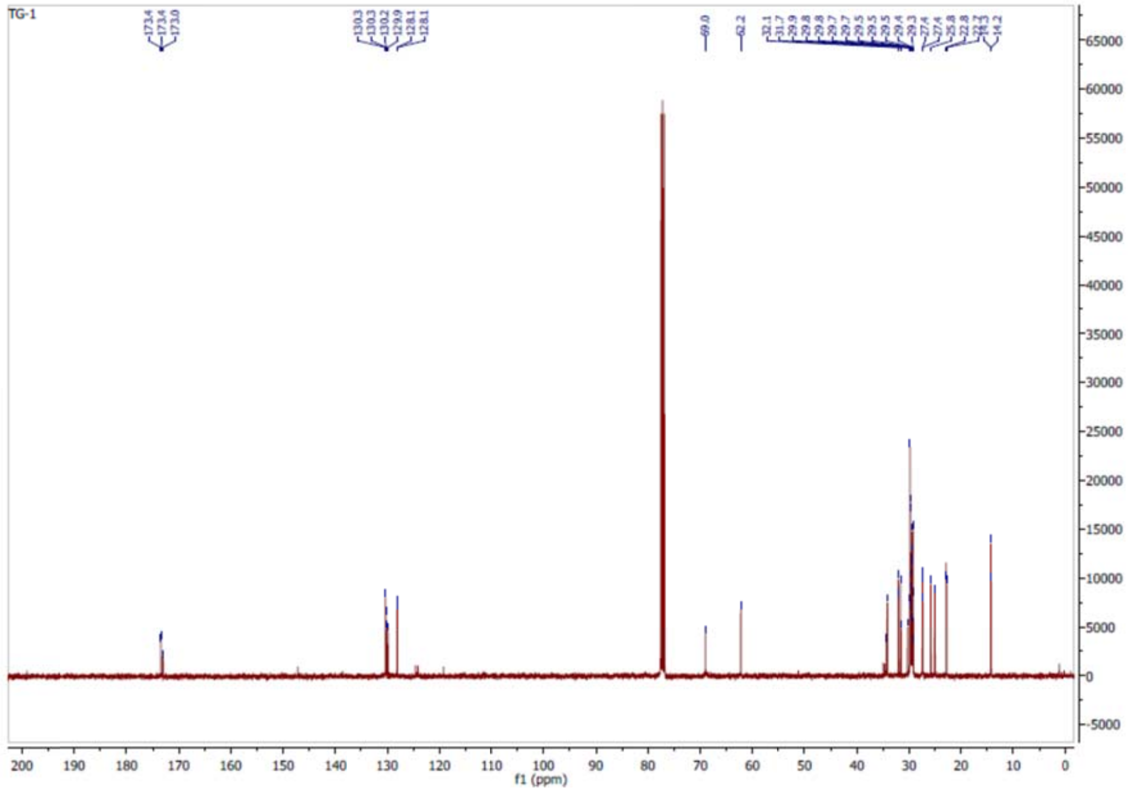

**TG (C16:0; C20:2, 11z,14z; C18:1, 9z),** ***(11Z,14Z)-1-(oleoyloxy)-3-(palmitoyloxy)propan-2-yl icosa-11,14-dienoate*****, TG2.**

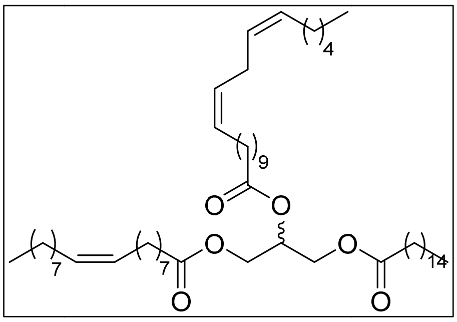

From 86 mg of **4** following the **procedure E** with (11Z,14Z)-icosa-11,14-dienoic acid 95 mg of **TG2** are obtained after purification by flash chromatography using Hexane/MTBE 3% as eluent (74% yield). $^{1}$H NMR (400 MHz, CDCl3) δ 5.40 – 5.30 (m, 6H), 5.26 (tt, *J* = 5.9, 4.3 Hz, 1H), 4.29 (dd, *J* = 11.9, 4.3 Hz, 2H), 4.14 (dd, *J* = 11.9, 6.0 Hz, 2H), 2.77 (t, *J* = 6.4 Hz, 2H), 2.31 (td, *J* = 7.6, 2.5 Hz, 6H), 2.10-1.96 (m, 8H), 1.66 – 1.57 (m, 6H), 1.39 – 1.19 (m, 62H), 0.88 (t, *J* = 6.8 Hz, 9H). $^{13}$C NMR (101 MHz, $CDCl_3$) δ 173.4, 173.4, 173.0, 130.3, 130.2, 130.1, 129.8, 128.1, 128.1, 69.0, 62.2, 34.3, 34.1, 34.1, 32.1, 32.0, 31.8, 29.9, 29.8, 29.8, 29.8, 29.8, 29.7, 29.6, 29.5, 29.5, 29.5, 29.4, 29.4, 29.4, 29.3, 29.2, 29.2, 27.4, 27.3, 25.8, 25.0, 25.0, 24.9, 22.8, 22.7, 14.2, 14.2. Anal. Calcd for C57H104O6 (885.45 g/mol): C, 77.32; H, 11.84%. Found: C, 77.38; H, 11.79%.

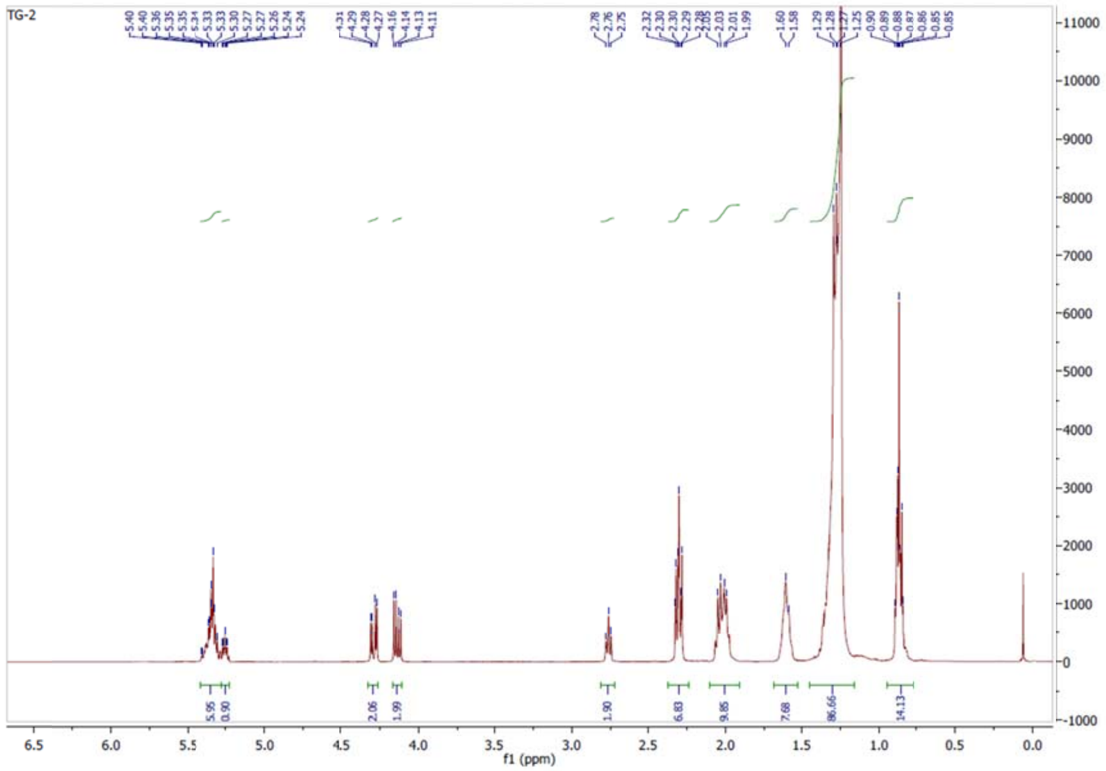
TG-2
f1 (ppm)

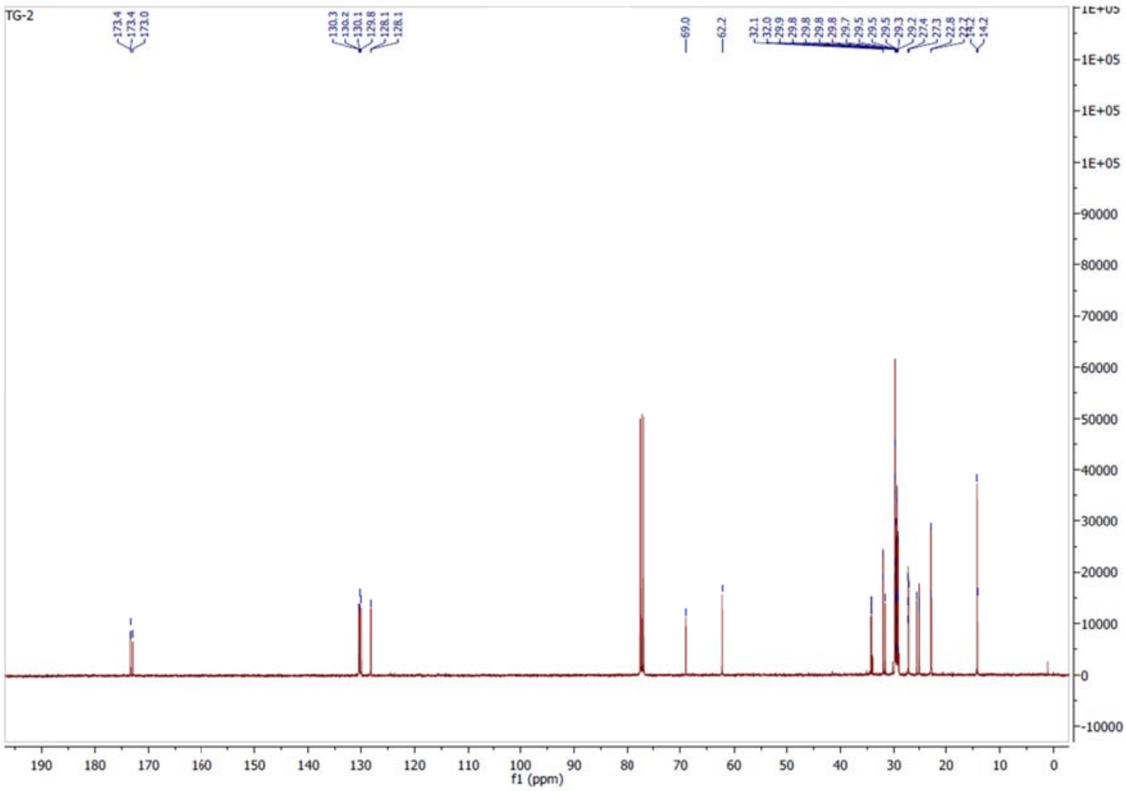
TG-2
f1 (ppm)

**TG (C16:0; C18:1, 9z; C18:1, 9z), (*Z)-3-(palmitoyloxy)propane-1,2-diyl dioleate*, TG3.**

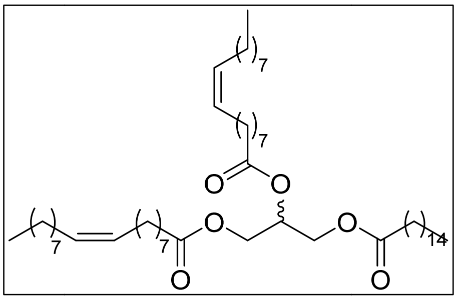

From 500 mg of **2** following the **procedure A** with oleoyl chloride 150 mg of **TG3** are obtained after purification by flash chromatography using Hexane/MTBE 3% as eluent (8% yield). In this step 540 mg of **4** is obtained as a colorless oil (60% yield) $^{1}$H NMR (400 MHz, $CDCl_3$) δ 5.39 – 5.30 (m, 4H), 5.26 (tt, *J* = 5.9, 4.3 Hz, 1H), 4.29 (dd, *J* = 11.9, 4.3 Hz, 2H), 4.14 (dd, *J* = 11.9, 6.0 Hz, 2H), 2.31 (td, *J* = 7.6, 2.5 Hz, 6H), 2.01 (q, *J* = 5.9, 5.3 Hz, 8H), 1.66 – 1.57 (m, 6H), 1.39 – 1.19 (m, 64H), 0.86-0.90 (m, 9H). $^{13}$C NMR (101 MHz, $CDCl_3$) δ 173.4, 173.4, 173.0, 130.3, 130.2, 130.1, 129.8, 69.0, 62.2, 34.3, 34.1, 34.1, 32.1, 32.0, 31.8, 29.9, 29.8, 29.8, 29.8, 29.8, 29.7, 29.6, 29.6, 29.5, 29.5, 29.4, 29.4, 29.4, 29.3, 29.2, 29.2, 27.4, 27.3, 25.8, 25.0, 25.0, 24.9, 22.8, 22.7, 14.3, 14.2. Anal. Calcd for C55H102O6 (859.42 g/mol): C, 76.87; H, 11.96%. Found: C, 76.94; H, 12.01%.

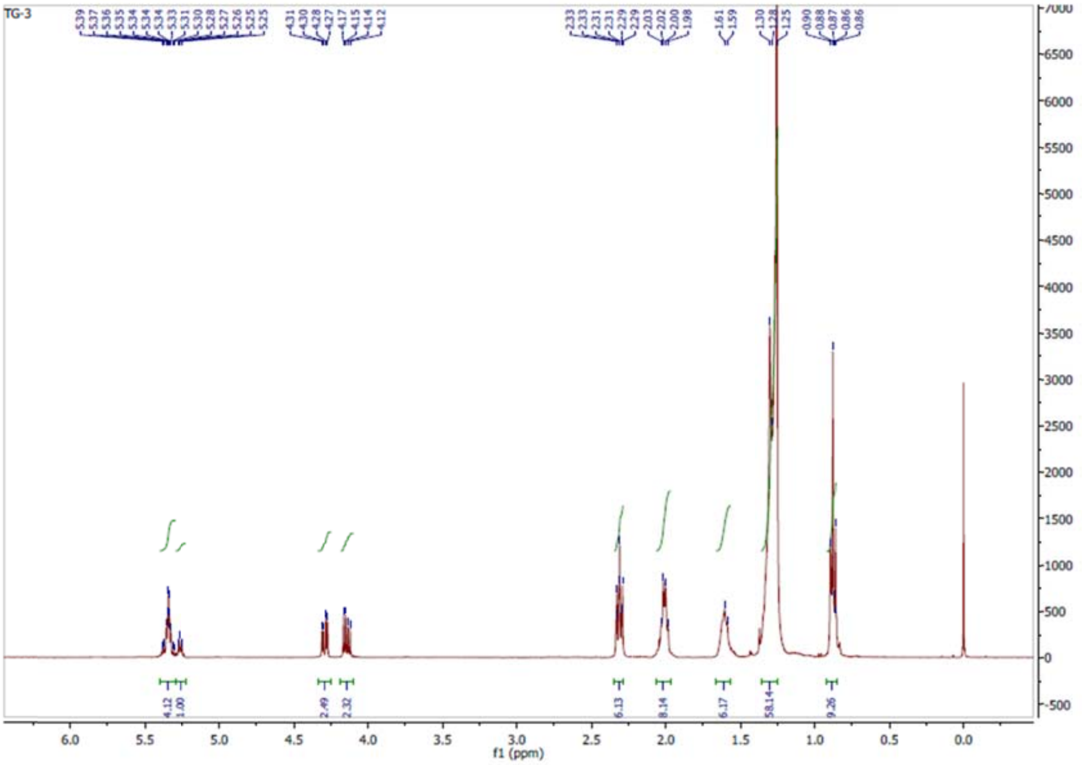

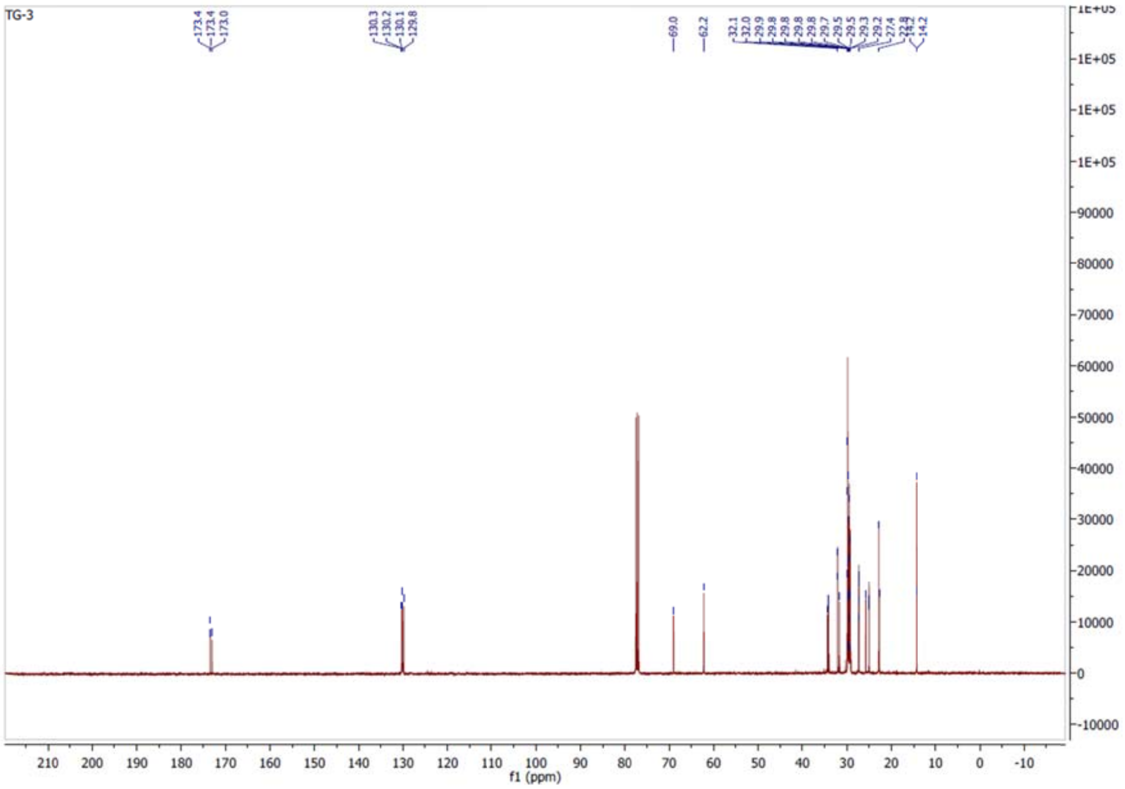

**TG (C18:0; C18:1, 9z; C18:2, 9z,12z),** ***(9Z,12Z)-2-(oleoyloxy)-3-(stearoyloxy)propyl octadeca-9,12-dienoate***, **TG4.**

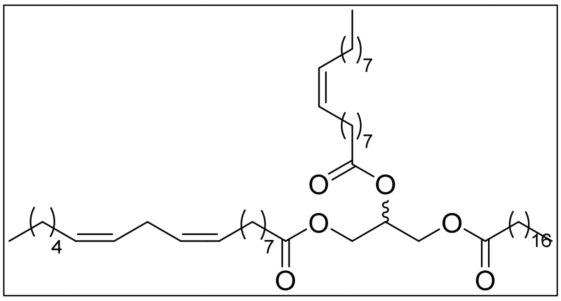

From 1 g of(±)-(2,2-dimethyl-1,3-dioxolan-4-yl)methanol, following the **procedure A** with stearic chloride, and **procedure B** to obtain **7**, which is transformed following the **procedure C**, followed by **procedure A** with oleoyl chloride to obtain 2.95 g of **8** (53%, 2 steps). 67 mg of **TG4** are obtained from 110 mg of **8** following the **procedure D** and **procedure A** with (*9Z*,*12Z*)-octadeca-9,12-dienoyl chloride after purification by flash chromatography using Hexane/MTBE 2% as eluent (62% yield, 2 steps). $^{1}$H NMR (400 MHz, $CDCl_3$) δ 5.41 – 5.29 (m, 6H), 5.26 (tt, *J* = 5.9, 4.6 Hz, 1H), 4.29 (dd, *J* = 11.9, 4.3 Hz, 2H), 4.14 (dd, *J* = 11.9, 6.0 Hz, 2H), 2.77 (t, *J* = 6.5 Hz, 2H), 2.31 (td, *J* = 7.5, 2.4 Hz, 6H), 2.07 – 1.98 (m, 8H), 1.65 – 1.57 (m, 4H), 1.38 – 1.22 (m, 64H), 0.90 – 0.86 (m, 9H). $^{13}$C NMR (101 MHz, $CDCl_3$) δ 173.4 173.4, 173.0, 130.3, 130.2, 130.1, 129.8, 128.2, 128.0, 69.0, 62.2, 34.3, 34.2, 34.1, 32.1, 32.0, 31.7, 29.9, 29.8, 29.8, 29.7, 29.7, 29.5, 29.5, 29.5, 29.4, 29.4, 29.4, 29.3, 29.3, 29.2, 29.2, 29.1, 27.3, 27.3, 25.8, 24.9, 24.9, 22.8, 22.7, 14.3, 14.2. Anal. Calcd for C57H104O6 (885.45 g/mol): C, 77.32; H, 11.84%. Found: C, 77.26; H, 11.79%.

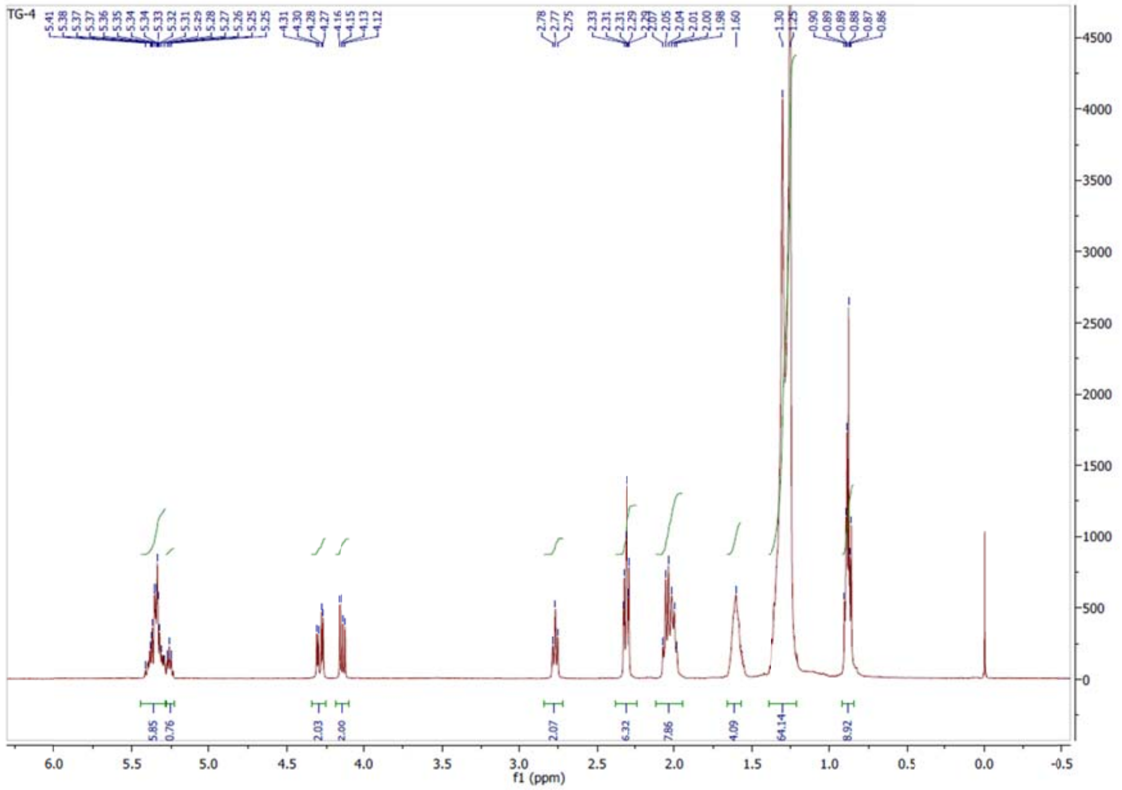
TG-4
f1 (ppm)

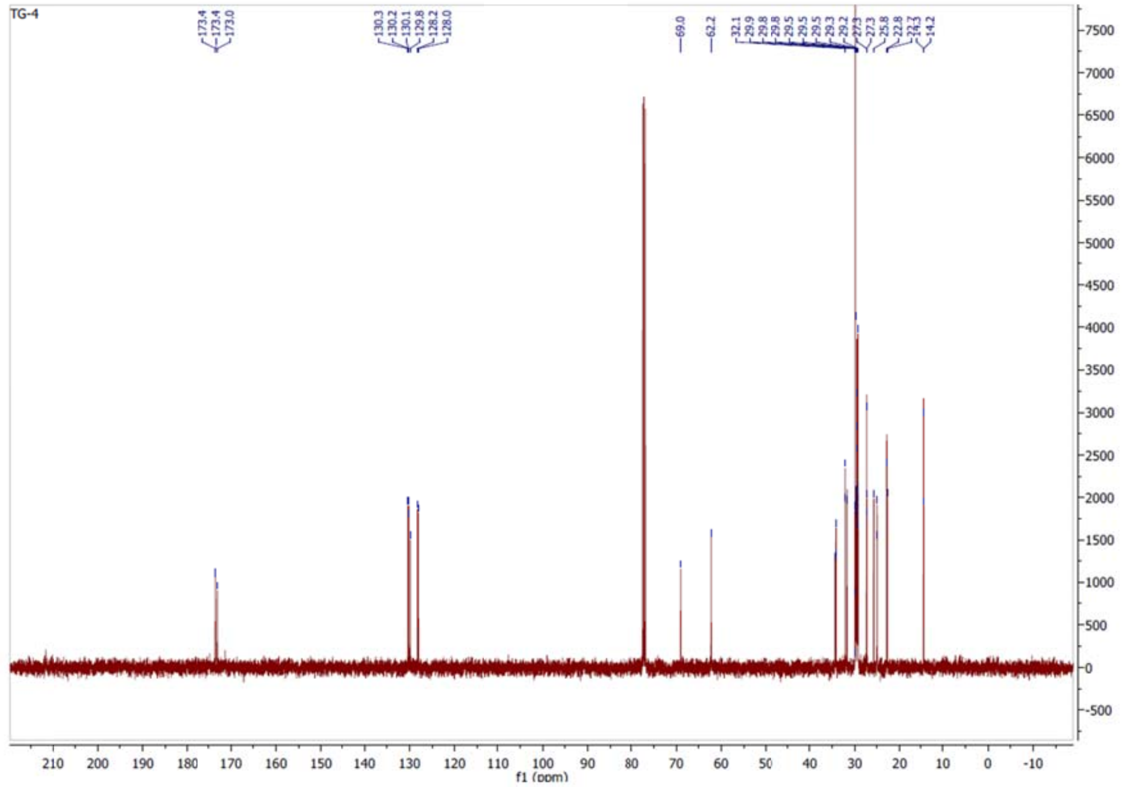
TG-4
f1 (ppm)

***2S*-TG (C16:0; C18:1, 9z; C20:2, 11z,14z), *(11Z,14Z)-(S)-2-(oleoyloxy)-3-(palmitoyloxy)propyl icosa-11,14-dienoate*, TG5.**

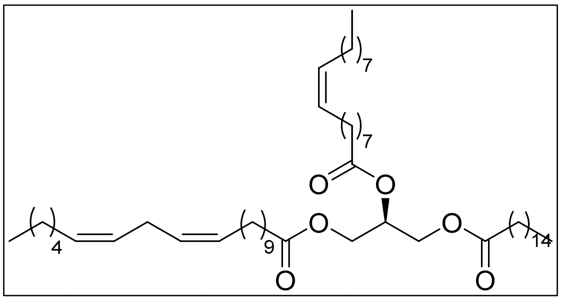

From 500 mg of (S)-(2,2-dimethyl-1,3-dioxolan-4-yl)methanol, following the **procedure A** with palmitoyl chloride, and **procedure B** to obtain **2a** with quantitative yield. **2a** is transformed following the **procedure C**, followed by **procedure A** with oleoyl chloride to obtain, after purification by flash chromatography using Hexane/MTBE 3% as eluent, 1.56 g of **3a** (58% yield, 4 steps). 88 mg of **TG5** is obtained from 76 mg of **3a** following the **procedure D** and **procedure E** with (11Z,14Z)-icosa-11,14-dienoic acid after purification by flash chromatography using Hexane/MTBE 3% as eluent (92% yield, 2 steps). $^{1}$H NMR (400 MHz, $CDCl_3$) δ 5.41 – 5.30 (m, 6H), 5.26 (td, *J* = 5.9, 3.0 Hz, 1H), 4.29 (dd, *J* = 11.9, 4.3 Hz, 2H), 4.14 (dd, *J* = 11.9, 5.9 Hz, 2H), 2.77 (t, *J* = 6.4 Hz, 2H), 2.31 (td, *J* = 7.6, 2.4 Hz, 6H), 2.10 – 1.94 (m, 8H), 1.67 – 1.57 (m, 6H), 1.41 – 1.18 (m, 62H), 0.93 – 0.84 (m, 9H). $^{13}$C NMR (101 MHz, $CDCl_3$) δ 173.4, 173.4, 173.0, 130.3, 130.2, 130.1, 129.8, 128.1, 128.0, 69.0, 62.2, 34.2, 32.0, 32.1, 31.7, 30.3, 29.9, 29.8, 29.8, 29.8, 29.7, 29.6, 29.6, 29.5, 29.5, 29.5, 29.4, 29.4, 29.4, 29.3, 29.2, 27.3, 27.3, 27.4, 27.3, 25.8, 25.0, 25.0, 22.8, 22.7, 14.3, 14.2. Anal. Calcd for C57H104O6 (885.45 g/mol): C, 77.32; H, 11.84%. Found: C, 77.27; H, 11.90%.

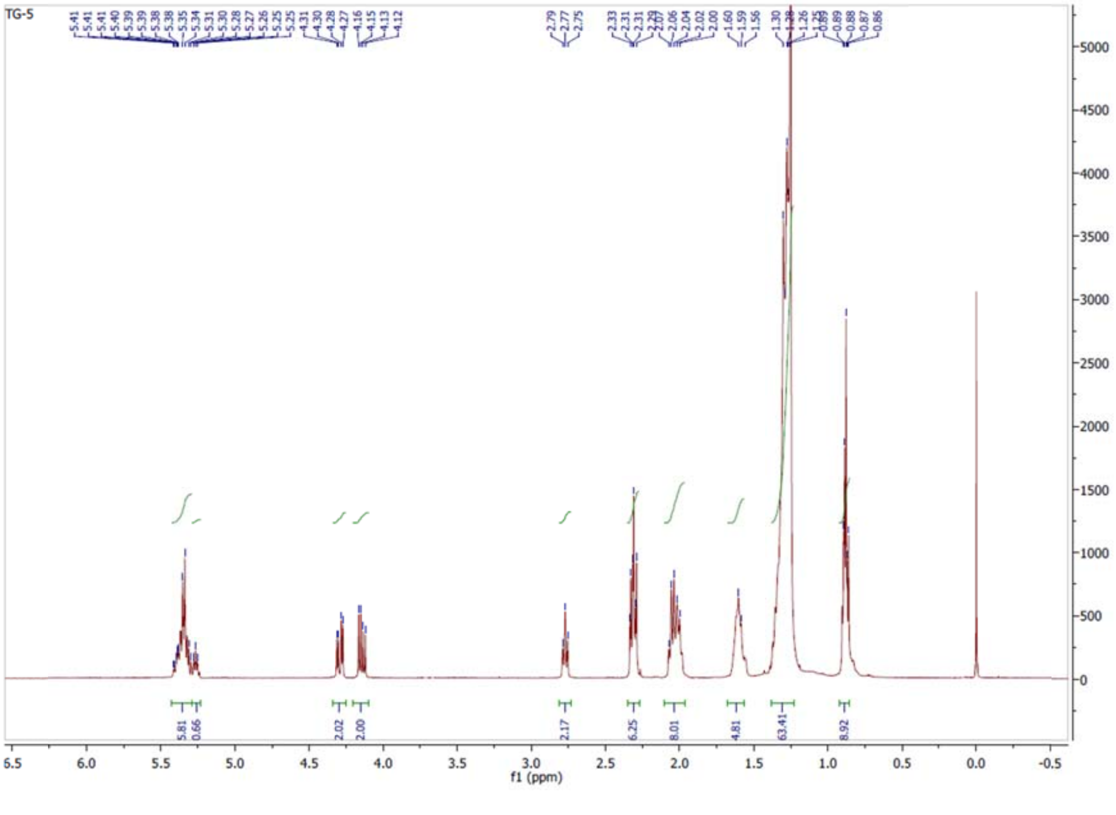

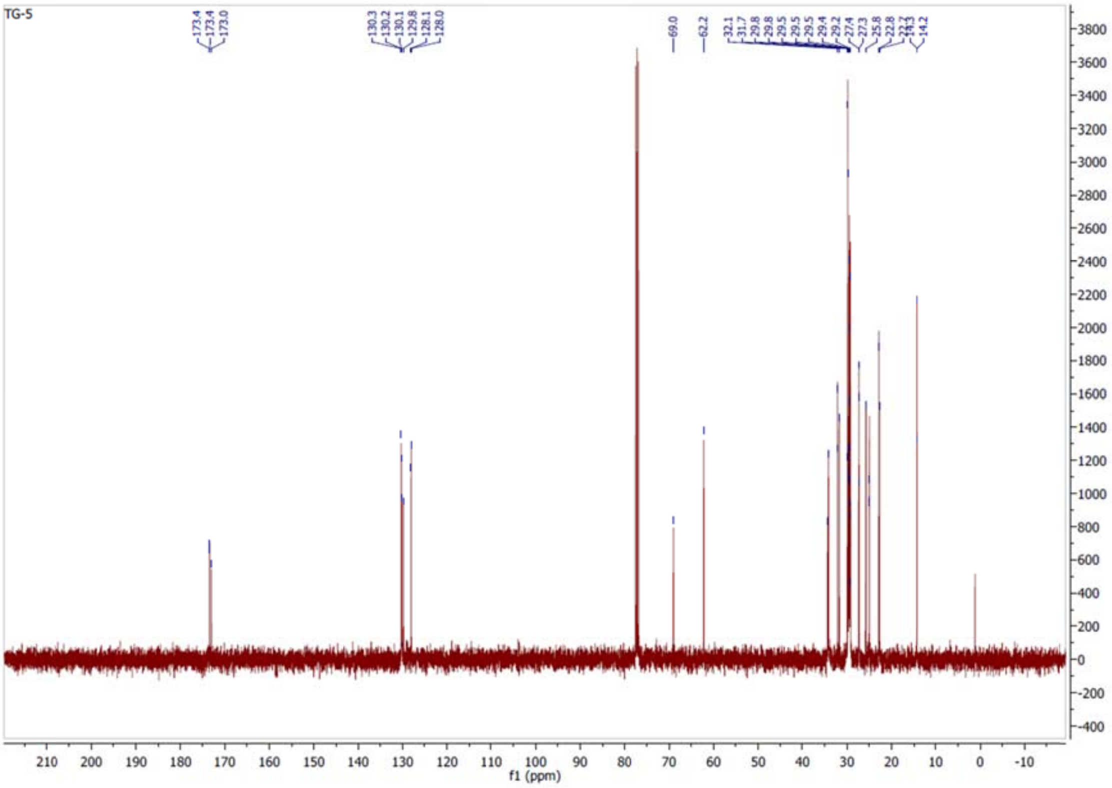

***2R*-TG (C16:0; C18:1, 9z; C20:2, 11z,14z), *(11Z,14Z)-(S)-2-(oleoyloxy)-3-(palmitoyloxy)propyl icosa-11,14-dienoate*, TG6.**

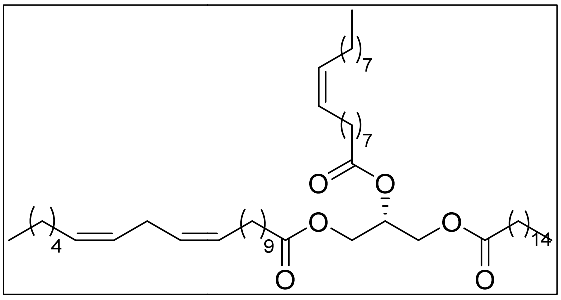

From 431 mg of (*R*)-(2,2-dimethyl-1,3-dioxolan-4-yl)methanol, following the **procedure A** with palmitoyl chloride, and **procedure B** to obtain **2b** with quantitative yield. **2b** is transformed following the **procedure C**, followed by **procedure A** with oleoyl chloride to obtain, after purification by flash chromatography using Hexane/MTBE 3% as eluent, 1.27 g of **3b** (55% yield, 4 steps). 77 mg of **TG6** is obtained from 76 mg of **3b** following the **procedure D** and **procedure E** with (11Z,14Z)-icosa-11,14-dienoic acid after purification by flash chromatography using Hexane/MTBE 3% as eluent (81% yield, 2 steps). $^1$H NMR (400 MHz, $CDCl_3$) δ 5.42 – 5.28 (m, 6H), 5.26 (td, *J* = 5.9, 3.0 Hz, 1H), 4.29 (dd, *J* = 11.9, 4.3 Hz, 2H), 4.14 (dd, *J* = 11.9, 5.9 Hz, 2H), 2.77 (t, *J* = 6.4 Hz, 2H), 2.31 (td, *J* = 7.6, 2.4 Hz, 6H), 2.10 – 1.94 (m, 8H), 1.67 – 1.57 (m, 6H), 1.41 – 1.18 (m, 62H), 0.93 – 0.84 (m, 9H). $^{13}$C NMR (101 MHz, $CDCl_3$) δ 173.4, 173.4, 173.0, 130.3, 130.2, 130.1, 129.8, 128.1, 128.0, 69.0, 62.2, 34.2, 32.1, 32.0, 31.7, 30.3, 29.9, 29.8, 29.8, 29.8, 29.7, 29.6, 29.6, 29.6, 29.5, 29.5, 29.4, 29.4, 29.4, 29.3, 29.2, 27.3, 27.3, 27.4, 27.3, 25.8, 25.0, 25.0, 22.8, 22.7, 14.3, 14.2. Anal. Calcd for C57H104O6 (885.45 g/mol): C, 77.32; H, 11.84%. Found: C, 77.36; H, 11.89%.

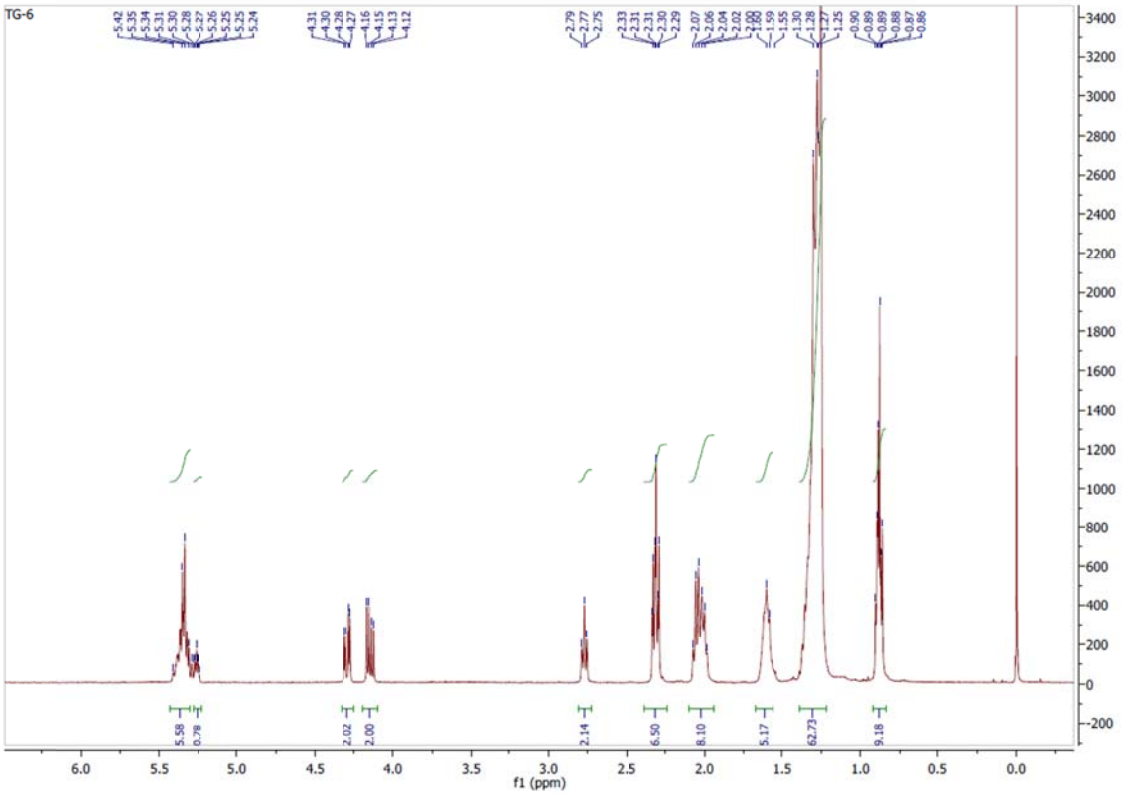

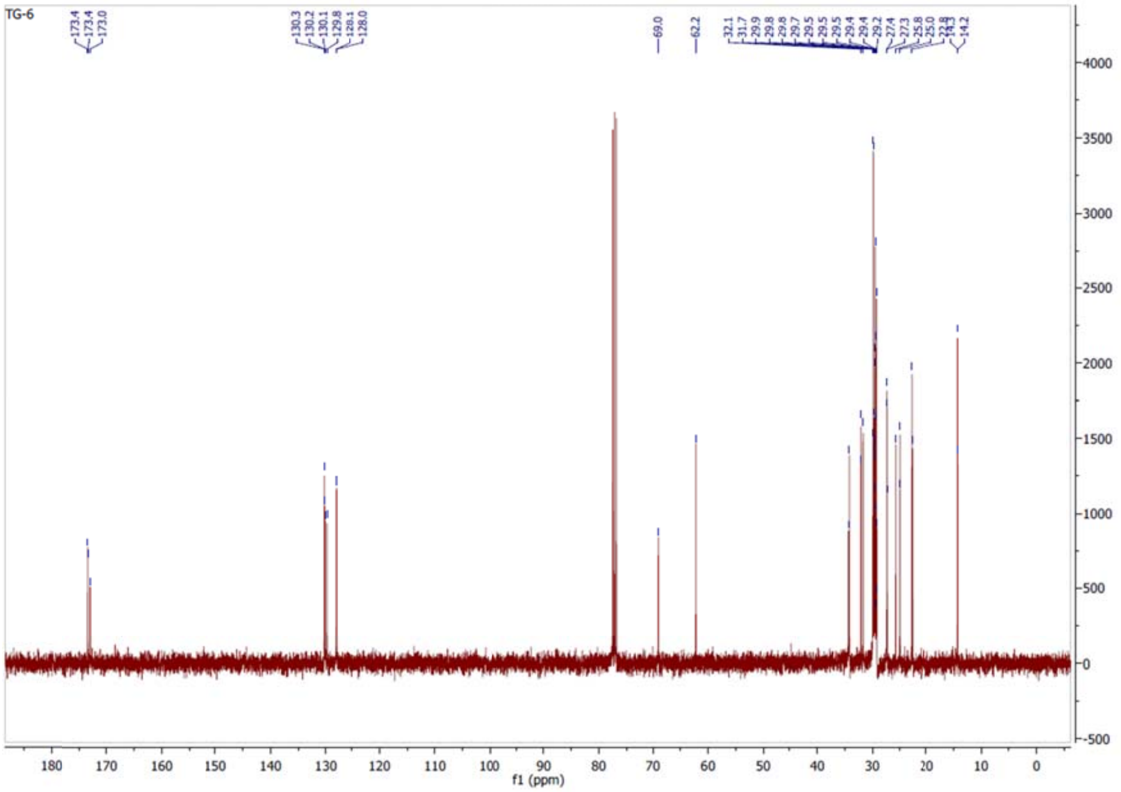

***2S*-TG (C20:2, 11z,14z; C16:0; C18:1, 9z), *(11Z,14Z)-(S)-3-(oleoyloxy)-2-(palmitoyloxy)propyl icosa-11,14-dienoate*, TG7.**

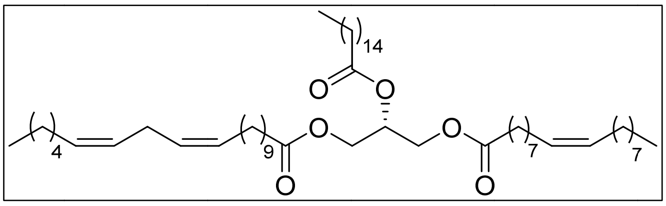

From 500 mg of (*R*)-(2,2-dimethyl-1,3-dioxolan-4-yl)methanol, following the **procedure A** with oleyl chloride, and **procedure B** to obtain **5a** with quantitative yield. **5a** is transformed following the **procedure C**, followed by **procedure A** with palmitoyl chloride to obtain, after purification by flash chromatography using Hexane/MTBE 1% as eluent, 1.43 g of **6a** (53% yield, 4 steps). 90 mg of **TG7** are obtained from 77 mg of **6a** following the **procedure D** and **procedure E** with (11Z,14Z)-icosa-11,14-dienoic acid after purification by flash chromatography using Hexane/MTBE 3% as eluent (95% yield, 2 steps). $^{1}$H NMR (400 MHz, $CDCl_3$) δ 5.41 – 5.29 (m, 6H), 5.27 (tt, *J* = 6.0, 4.4 Hz, 1H), 4.29 (dd, *J* = 11.9, 4.3 Hz, 2H), 4.14 (dd, *J* = 11.9, 6.0 Hz, 2H), 2.77 (t, *J* = 6.5 Hz, 2H), 2.31 (td, *J* = 7.5, 2.4 Hz, 6H), 2.10 – 1.96 (m, 8H), 1.66 – 1.56 (m, 6H), 1.37 – 1.21 (m, 62H), 0.93 – 0.84 (m, 9H). $^{13}$C NMR (101 MHz, $CDCl_3$) δ 173.3, 173.3, 172.9, 130.3, 130.2, 130.1, 129.8, 128.1, 128.0, 69.0, 62.2, 34.3, 34.1, 32.1, 32.0, 31.7, 29.9, 29.8, 29.8, 29.7, 29.6, 29.5, 29.5, 29.4, 29.4, 29.4, 29.3, 29.2, 29.2, 27.3, 27.3, 27.2, 25.7, 25.0, 24.9, 22.8, 22.7, 14.2, 14.2. Anal. Calcd for C57H104O6 (885,43 g/mol): C, 77.32; H, 11.84% Found: C, 77.13; H, 11.75%.

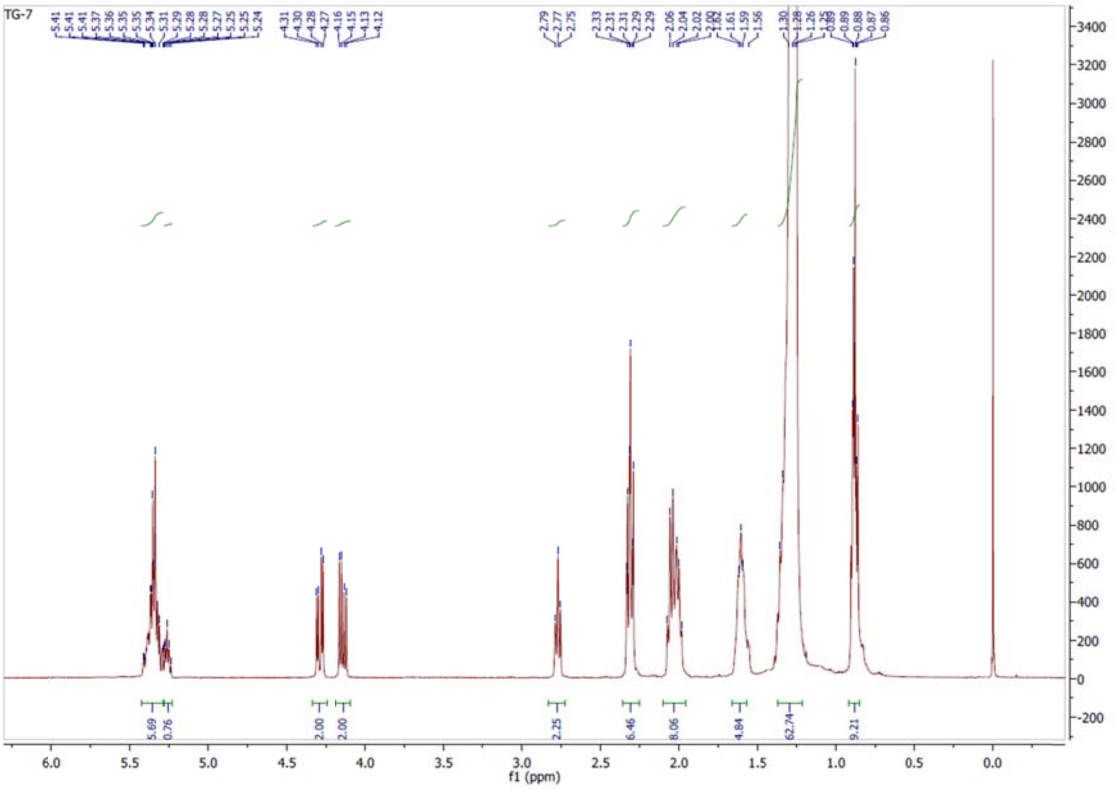

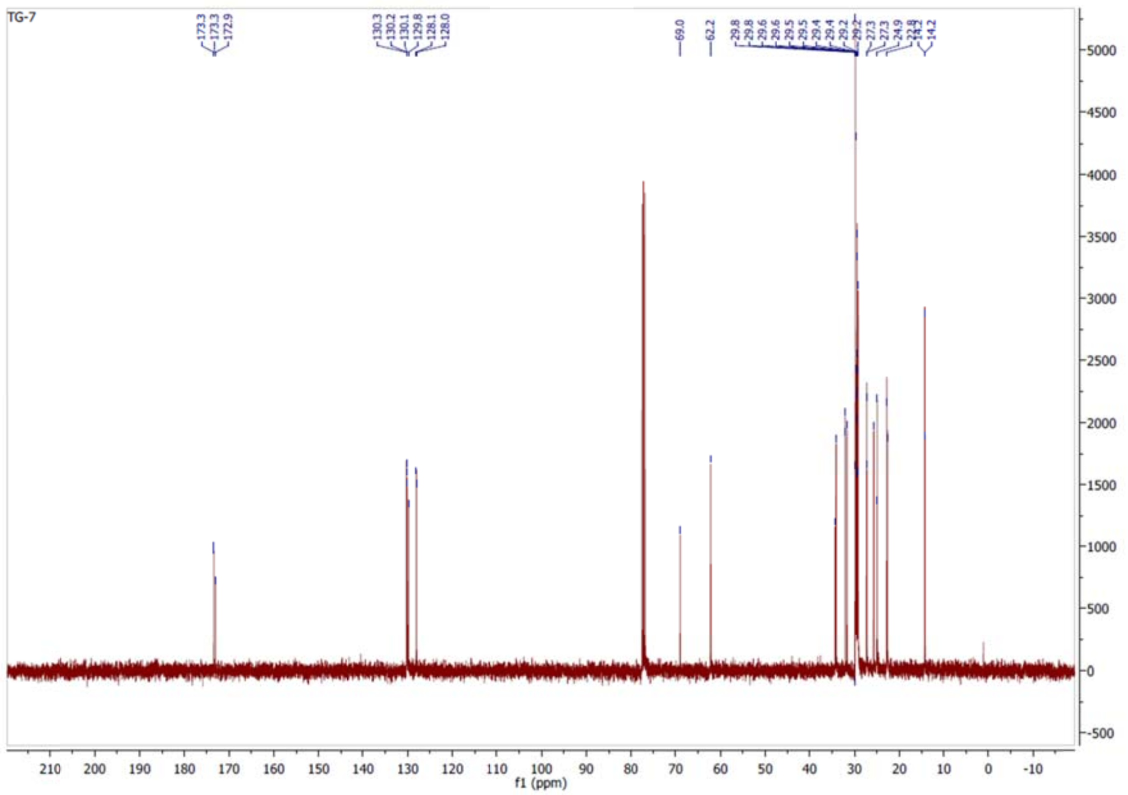

***2R*-TG (C20:2, 11z,14z; C16:0; C18:1, 9z), *(11Z,14Z)-(R)-3-(oleoyloxy)-2-(palmitoyloxy)propyl icosa-11,14-dienoate*, TG8.**

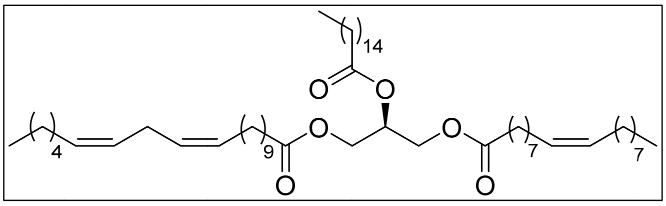

From 500 mg of (*S*)-(2,2-dimethyl-1,3-dioxolan-4-yl)methanol, following the **procedure A** with oleyl chloride, and **procedure B** to obtain **5b** with quantitative yield. **5b** is transformed following the **procedure C**, followed by **procedure A** with palmitoyl chloride to obtain, after purification by flash chromatography using Hexane/MTBE 1% as eluent, 1.16 g of **6b** (43% yield, 4 steps). 83 mg of **TG8** are obtained from 77 mg of **6b** following the **procedure D** and **procedure E** with (11Z,14Z)-icosa-11,14-dienoic acid after purification by flash chromatography using Hexane/MTBE 3% as eluent (87% yield, 2 steps). $^{1}$H NMR (400 MHz, $CDCl_3$) δ 5.42 – 5.29 (m, 6H), 5.27 (tt, *J* = 5.9, 4.6 Hz, 1H), 4.29 (dd, *J* = 11.9, 4.3 Hz, 2H), 4.14 (dd, *J* = 11.9, 6.0 Hz, 2H), 2.77 (t, *J* = 6.5 Hz, 2H), 2.31 (td, *J* = 7.5, 2.4 Hz, 6H), 2.10 – 1.95 (m, 8H), 1.70 – 1.51 (m, 6H), 1.40 – 1.22 (m, 62H), 0.94 – 0.84 (m, 9H). $^{13}$C NMR (101 MHz, $CDCl_3$) δ 173.3, 173.3, 173.0, 130.3, 130.2, 130.1, 129.8, 128.1, 128.0, 69.0, 62.2, 34.3, 34.2, 32.1, 32.0, 31.7, 29.9, 29.8, 29.8, 29.7, 29.7, 29.6, 29.5, 29.5, 29.4, 29.4, 29.4, 29.4, 29.3, 29.2, 29.2, 27.3, 27.3, 27.2, 25.7, 25.0, 24.9, 22.8, 22.7, 14.2, 14.2. Calcd for C57H104O6 (885,43 g/mol): C, 77.32; H, 11.84. Found: C, 77.22; H, 11.77%.

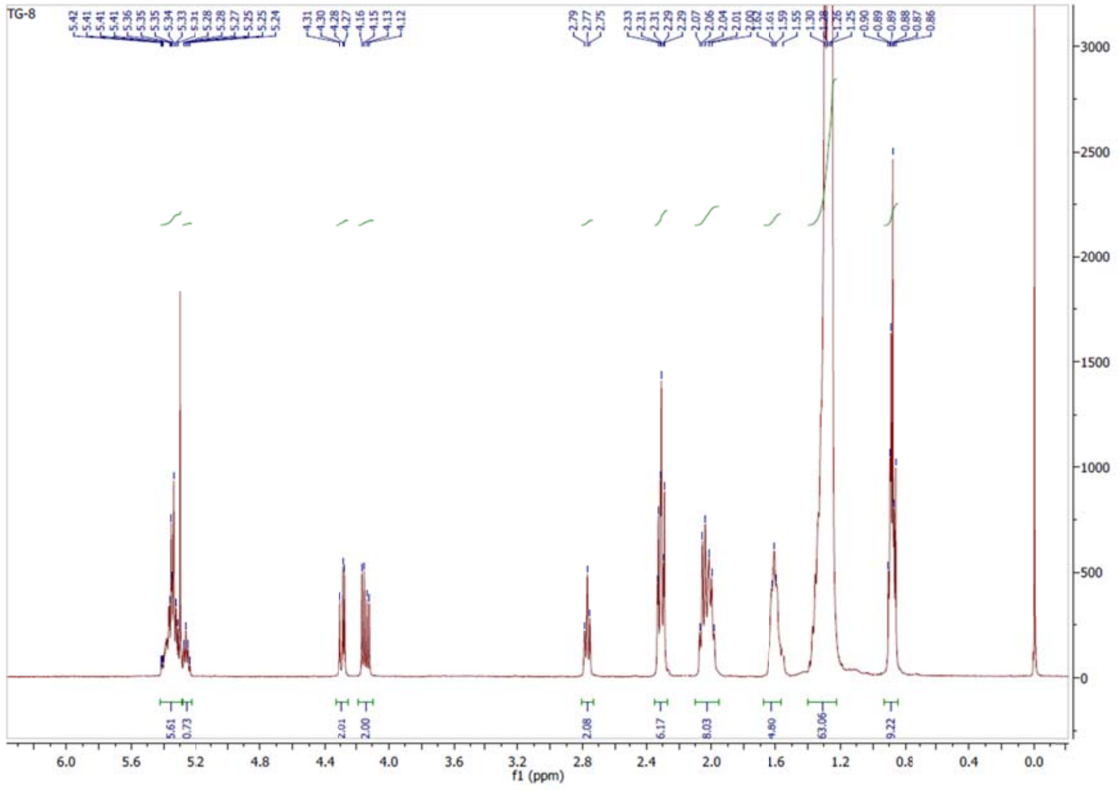
TG-8
f1 (ppm)

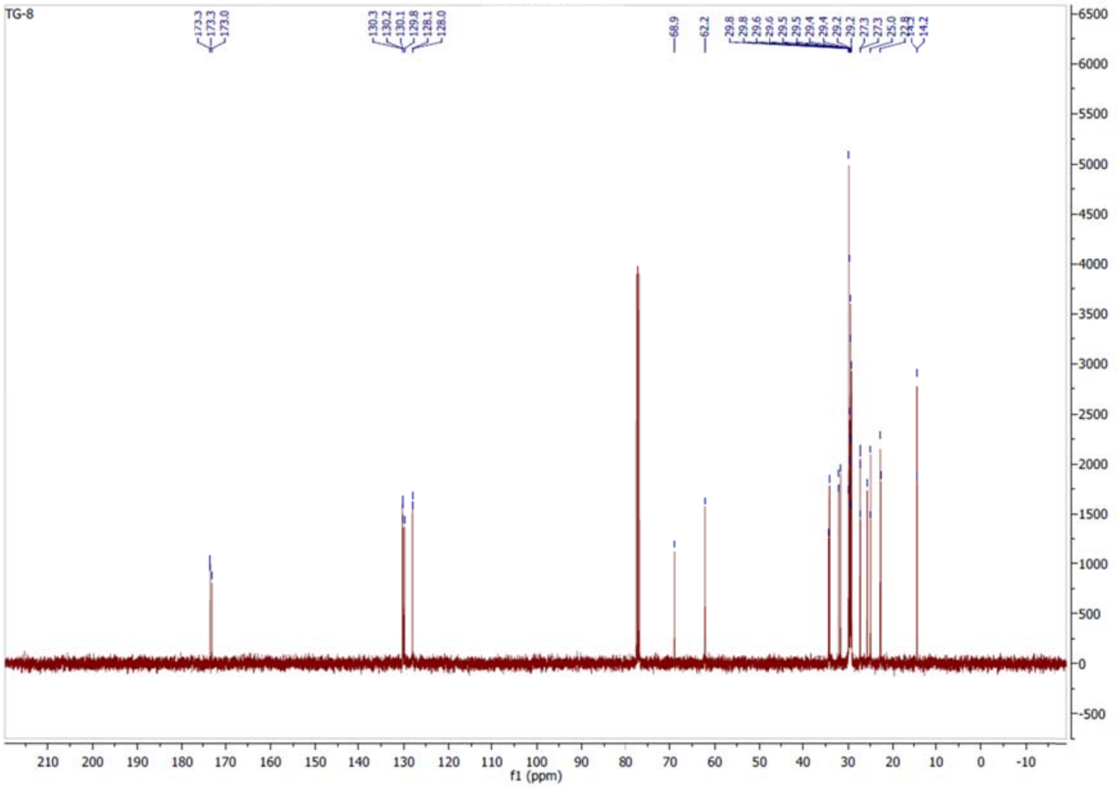
TG-8
f1 (ppm)

***2S*-TG (C18:2, 9z,12z; C16:0; C18:1, 9z), *(9Z,12Z)-(S)-3-(oleoyloxy)-2-(palmitoyloxy)propyl octadeca-9,12-dienoate*, TG9.**

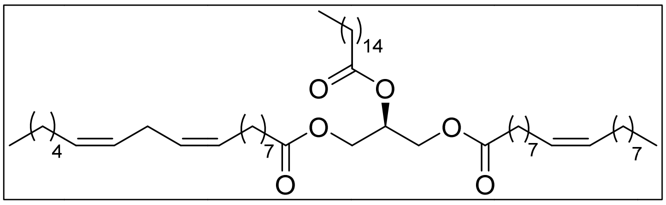

From 114 mg of **6a** following the **procedure D** and **procedure A** with linoleyl chloride after purification by flash chromatography using Hexane/MTBE 3% as eluent 104 mg of **TG9** are obtained (76% yield, 2 steps). $^{1}$H NMR (400 MHz, $CDCl_3$) δ 5.42 – 5.29 (m, 6H), 5.29 – 5.23 (m, 1H), 4.29 (dd, *J* = 11.9, 4.3 Hz, 2H), 4.14 (dd, *J* = 11.9, 6.0 Hz, 2H), 2.77 (t, *J* = 6.6 Hz, 2H), 2.31 (td, *J* = 7.5, 2.1 Hz, 6H), 2.10 – 1.96 (m, 8H), 1.67 – 1.56 (m, 4H), 1.41 – 1.19 (m, 60H), 0.95 – 0.82 (m, 9H). $^{13}$C NMR (101 MHz, $CDCl_3$) δ 173.3, 173.3, 172.9, 130.2, 130.1, 130.0, 129.7, 128.1, 127.9, 68.9, 62.1, 34.2, 34.1, 34.0, 32.0, 31.9, 31.5, 29.8, 29.7, 29.6, 29.6, 29.5, 29.5, 29.3, 29.3, 29.3, 29.3, 29.2, 29.1, 29.1, 27.2, 27.1, 25.6, 24.9, 24.8, 22.7, 22.6, 14.2, 14.1 Anal. Calcd for C55H100O6; C, 77.05; H, 11.76; (857,38 g/mol): Found: C, 76.85; H, 11.59%.

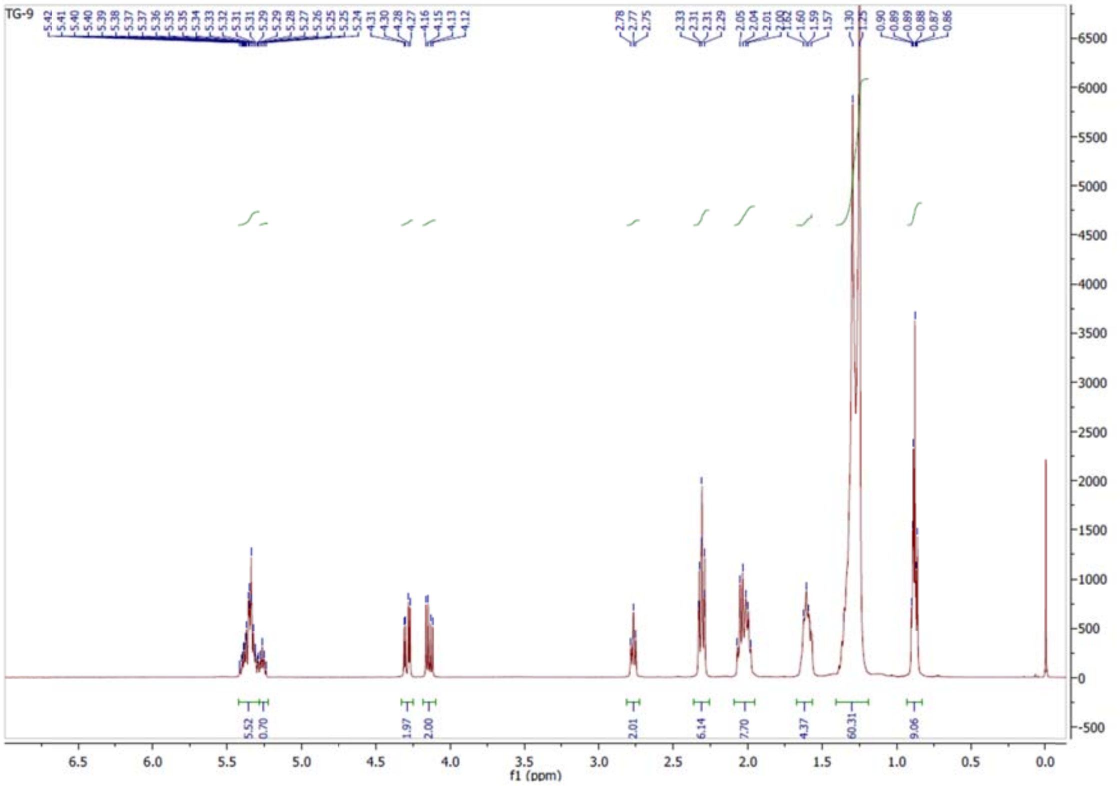

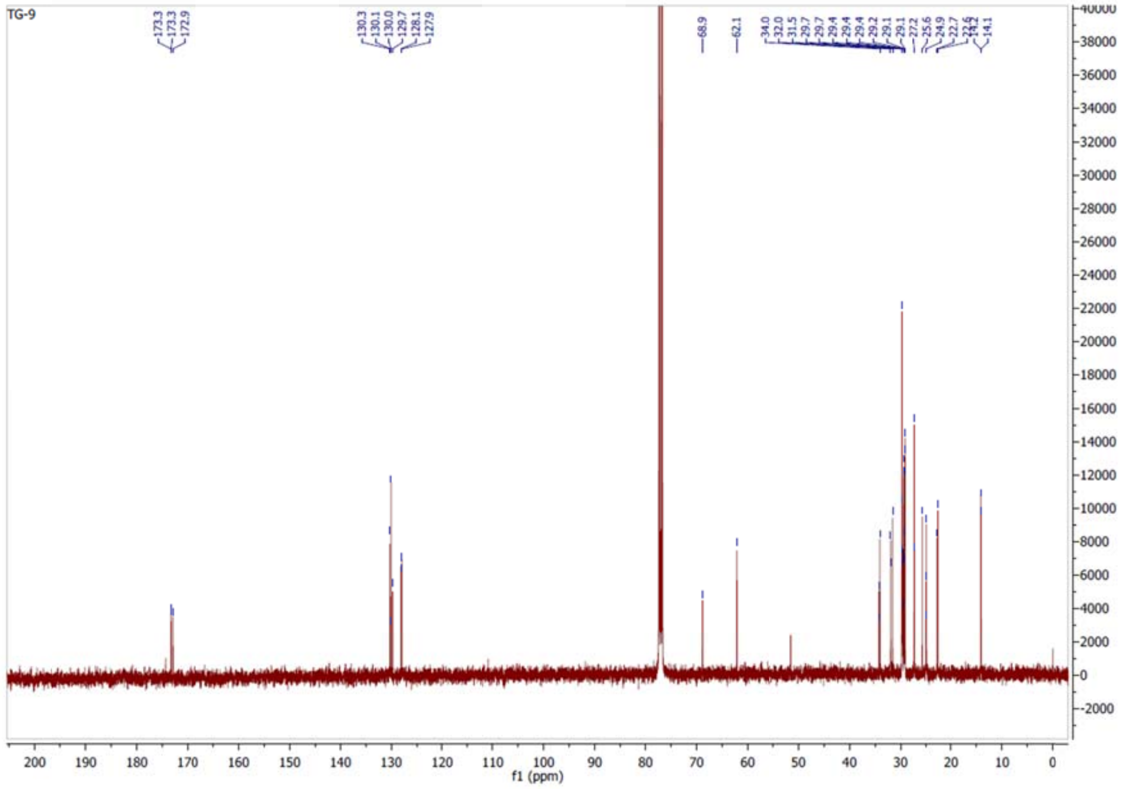

***2R*-TG (C18:2, 9z,12z; C16:0; C18:1, 9z), *(9Z,12Z)-(R)-3-(oleoyloxy)-2-(palmitoyloxy)propyl octadeca-9,12-dienoate*, TG10.**

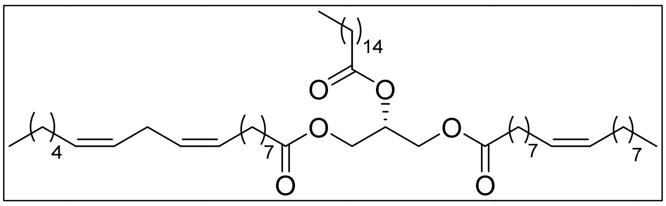

From 78 mg of **6b** following the **procedure D** and **procedure A** with linoleyl chloride after purification by flash chromatography using Hexane/MTBE 3% as eluent 94 mg of **TG10** are obtained (100% yield, 2 steps). $^{1}$H NMR (400 MHz, $CDCl_3$) δ 5.42 – 5.29 (m, 6H), 5.29 – 5.23 (m, 1H), 4.29 (dd, *J* = 11.9, 4.3 Hz, 2H), 4.14 (dd, *J* = 11.9, 6.0 Hz, 2H), 2.77 (t, *J* = 6.4 Hz, 2H), 2.31 (td, *J* = 7.5, 2.1 Hz, 6H), 2.09 – 1.96 (m, 8H), 1.66 – 1.58 (m, 4H), 1.45 – 1.17 (m, 60H), 0.94 – 0.84 (m, 9H). $^{13}$C NMR (101 MHz, $CDCl_3$) δ 173.3, 173.3, 172.9, 130.2, 130.1, 130.0, 129.7, 128.1, 127.9, 68.9, 62.1, 34.2, 34.1, 34.0, 32.0, 31.9, 31.5, 29.8, 29.7, 29.6, 29.6, 29.5, 29.5, 29.3, 29.3, 29.3, 29.3, 29.2, 29.1, 29.1, 27.2, 27.1, 25.6, 24.9, 24.8, 22.7, 22.6, 14.2, 14.1 Anal. Calcd for C55H100O6; C, 77.05; H, 11.76; (857,38 g/mol): Found: C, 76.92; H, 11.64%.

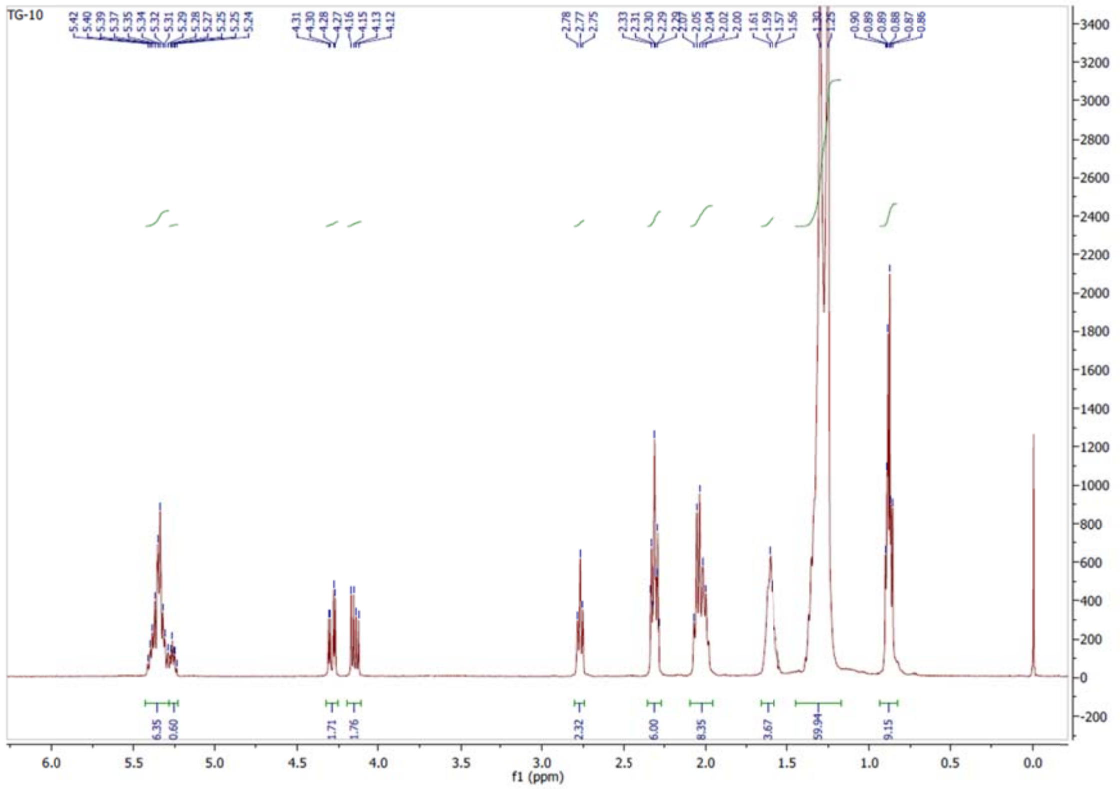
TG-10
f1 (ppm)

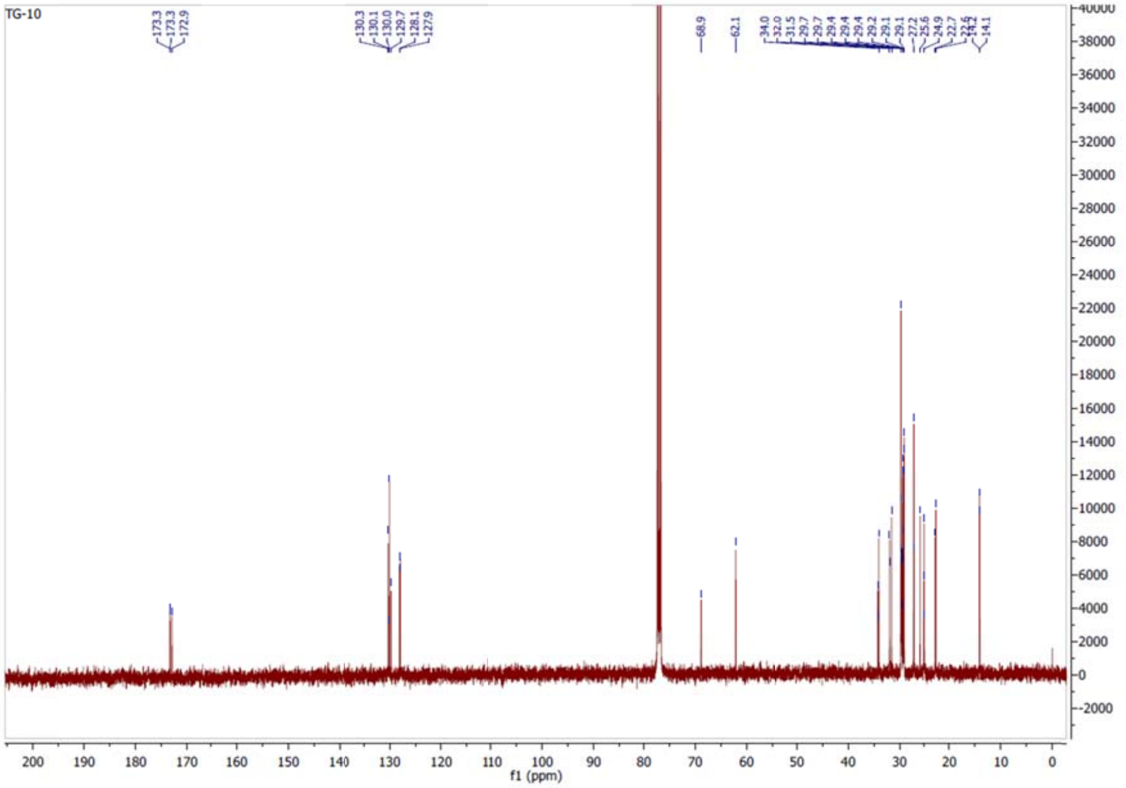
TG-10
f1 (ppm)

#### 3. Growth modulation activity of synthetic triglycerides

Cell lines A549 (blue) and NL20 (red) were exposed to different concentration of the indicated triglycerides (TG) for 5 days, and the optical density (O.D.) was measured.

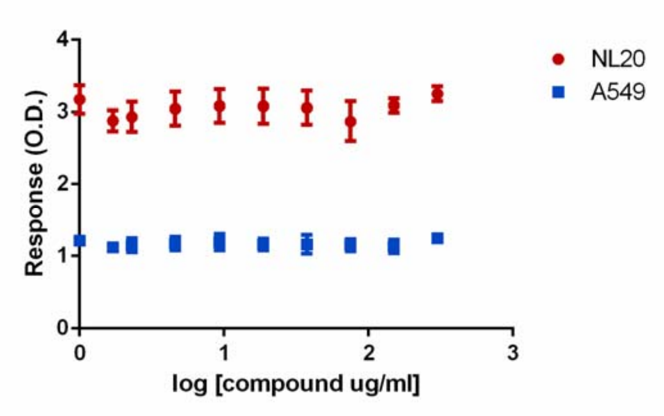

**TG1**

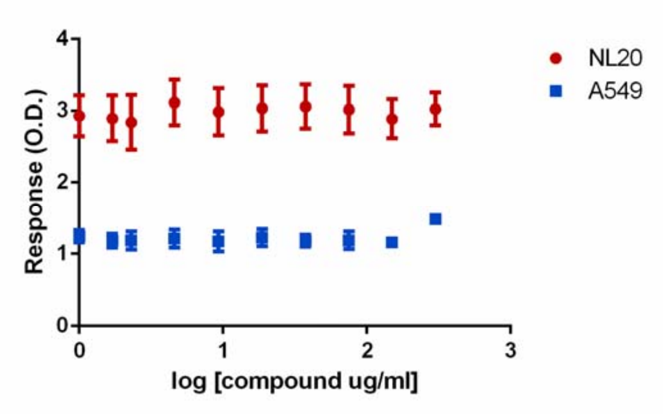

**TG2**

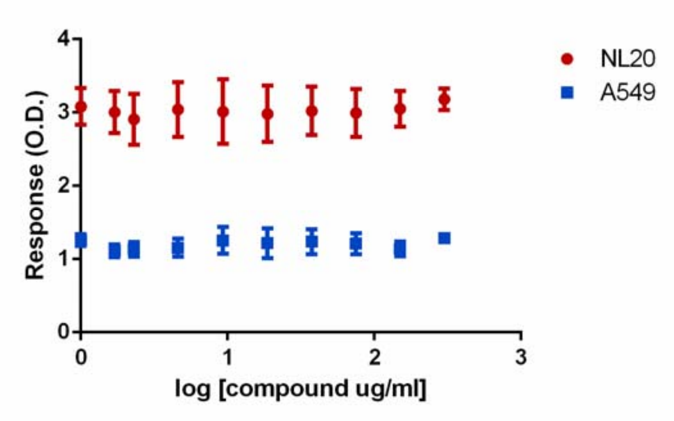

**TG3**

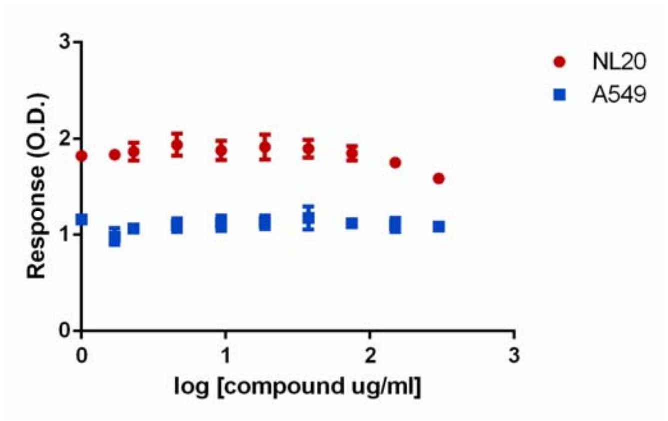
NL20
A549
Response (O.D.)
log [compound ug/ml]

**TG4**

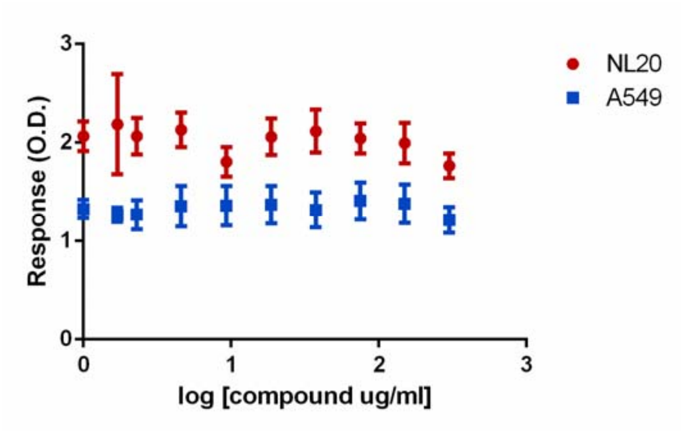
NL20
A549
Response (O.D.)
log [compound ug/ml]

**TG5**

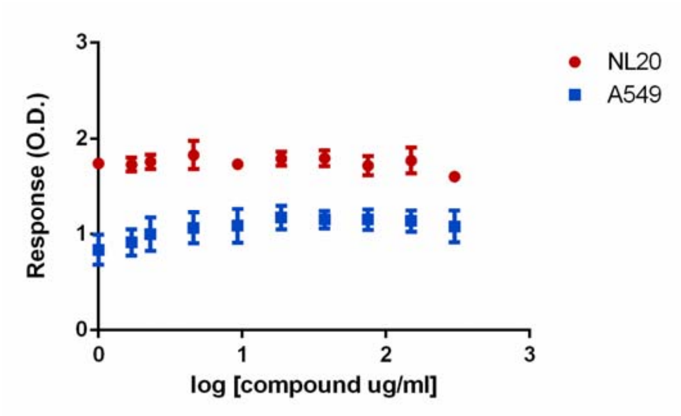
NL20
A549
Response (O.D.)
log [compound ug/ml]

**TG6**

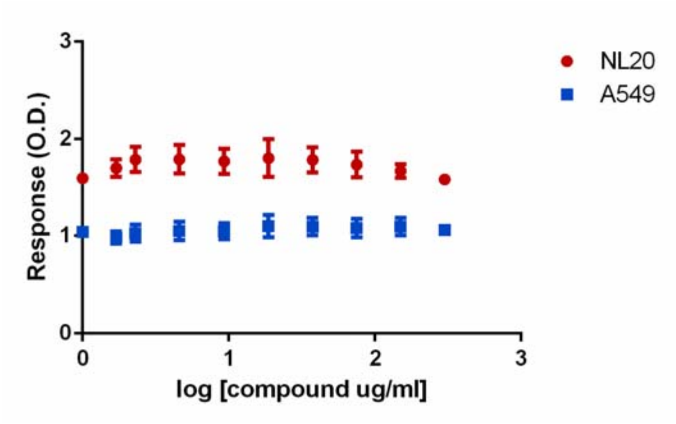

**TG7**

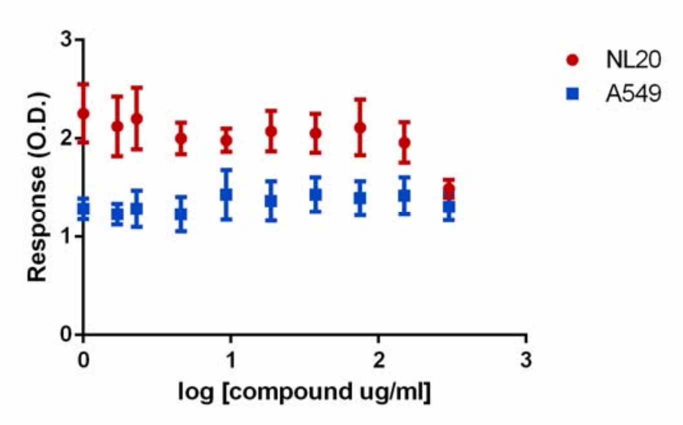

**TG8**

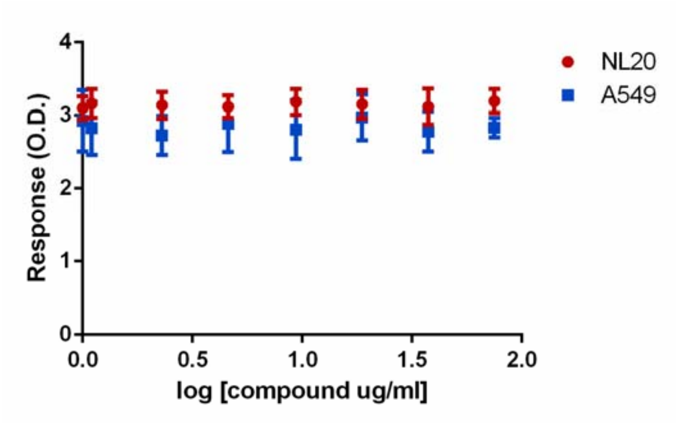

**TG9**

Results for **TG10** are shown in Figure 4 (main manuscript).

**4. Comparison of $^{13}$C-NMR spectra of aliphatic carbons between the synthetic triglyceride TG10 (red) and the natural extract (green)**

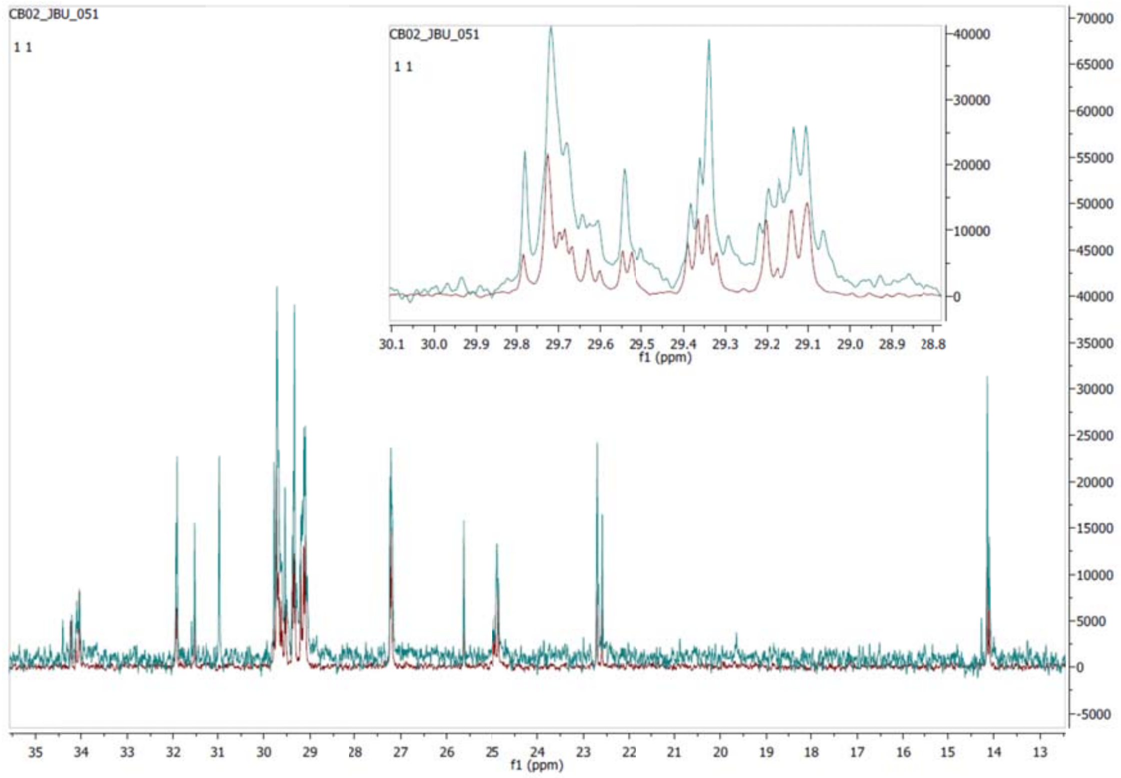

### 5. Dataset for xenograft study

Mice were injected with PBS as a control (AZ30-AZ45) or with Macrocybin (AZ26-AZ43) at the indicated days (d) after A549 tumor cell injection. Values represent tumor volume in mm3. Mouse AZ40 had to be sacrificed early because of a corneal ulcer.

| Mouse codes | d14 | d15 | d18 | d20 | d22 | d25 |
|---|---|---|---|---|---|---|
| AZ30 | 95,57 | 75,83 | 91,54 | 83,77 | 139,81 | 279,34 |
| AZ31 | 50,77 | 61,61 | 95,04 | 125,98 | 173,27 | 186,49 |
| AZ32 | 55,87 | 84,48 | 125,79 | 109,28 | 146,66 | 181,39 |
| AZ33 | 39,36 | 58,94 | 88,63 | 111,16 | 192,63 | 182,12 |
| AZ34 | 24,02 | 11,07 | 32,97 | 167,27 | 198,16 | 235,59 |
| AZ35 | 12,75 | 34,17 | 73,38 | 96,70 | 131,35 | 211,13 |
| AZ36 | 55,62 | 91,79 | 92,61 | 132,30 | 116,04 | 284,34 |
| AZ37 | 36,50 | 42,95 | 80,62 | 98,38 | 111,04 | 173,21 |
| AZ44 | 10,05 | 23,72 | 36,62 | 50,46 | 80,18 | 127,93 |
| AZ45 | 35,12 | 63,25 | 98,20 | 87,02 | 131,73 | 149,91 |
| AZ26 | 53,68 | 99,04 | 123,99 | 191,01 | 221,17 | 267,96 |
| AZ27 | 77,90 | 64,15 | 92,66 | 105,84 | 153,49 | 247,92 |
| AZ28 | 31,13 | 27,73 | 27,68 | 24,57 | 44,92 | 44,96 |
| AZ29 | 48,45 | 32,06 | 59,59 | 70,39 | 55,18 | 73,86 |
| AZ38 | 42,69 | 49,37 | 104,25 | 105,66 | 130,25 | 195,44 |
| AZ39 | 13,46 | 17,98 | 25,03 | 26,72 | 37,21 | 47,87 |
| AZ41 | 42,39 | 44,37 | 60,34 | 94,85 | 150,74 | 170,39 |
| AZ42 | 20,66 | 21,67 | 28,82 | 23,41 | 38,40 | 98,73 |
| AZ43 | 23,07 | 38,30 | 60,49 | 81,87 | 81,90 | 159,54 |

| d27 | d29 | d32 | d34 | d36 | d39 | d41 |
|---|---|---|---|---|---|---|
| 369,74 | 389,34 | 743,12 | 902,34 | 762,19 | 768,72 | 857,36 |
| 325,34 | 330,22 | 462,28 | 527,49 | 584,38 | 666,55 | 658,87 |
| 269,18 | 267,80 | 334,16 | 429,24 | 486,11 | 651,23 | 560,13 |
| 379,01 | 509,11 | 724,79 | 818,50 | 1108,49 | 1019,83 | 1140,44 |
| 259,53 | 286,32 | 644,73 | 830,32 | 839,56 | 858,94 | 1147,14 |
| 282,99 | 403,07 | 435,30 | 488,34 | 547,00 | 764,69 | 1072,99 |
| 177,84 | 483,79 | 908,66 | 913,59 | 1059,99 | 1032,26 | 1028,44 |
| 267,51 | 303,01 | 375,60 | 481,48 | 533,45 | 549,21 | 552,92 |
| 136,89 | 181,23 | 213,25 | 274,20 | 282,35 | 272,90 | 297,05 |
| 160,81 | 183,79 | 239,27 | 255,49 | 288,24 | 340,70 | 375,91 |
| 424,58 | 487,64 | 580,55 | 660,80 | 689,36 | 845,59 | 857,00 |
| 297,63 | 508,09 | 694,83 | 573,68 | 742,22 | 981,39 | 1087,49 |
| 39,94 | 72,26 | 99,72 | 134,73 | 97,36 | 185,48 | 182,20 |
| 117,74 | 120,27 | 131,38 | 171,26 | 157,68 | 209,00 | 183,03 |
| 211,87 | 228,24 | 310,53 | 307,82 | 292,13 | 376,58 | 428,24 |
| 45,51 | 69,49 | 132,41 | 112,55 | 90,93 | 150,94 | 153,75 |
| 207,31 | 196,98 | 254,95 | 282,18 | 412,03 | 624,50 | 540,00 |
| 80,90 | 104,52 | 190,06 | 196,01 | 230,97 | 301,75 | 333,86 |
| 179,28 | 209,38 | 233,72 | 306,08 | 371,39 | 500,67 | 494,62 |

| d43 | d46 | d48 | d50 | d53 | d55 |
|---|---|---|---|---|---|
| 1052,36 | 1099,10 | 1022,14 | 1303,67 | 1417,85 | 1655,35 |
| 606,17 | 546,98 | 694,16 | 1250,02 | 1027,84 | 1027,35 |
| 671,60 | 729,03 | 721,88 | 924,49 | | |
| 1443,88 | 1381,97 | 1292,36 | 1765,38 | 2082,13 | 2417,64 |
| 1192,24 | 1548,99 | 1636,23 | 1786,41 | 1924,15 | 2123,91 |
| 889,24 | 1020,19 | 977,39 | 1438,55 | 1485,81 | 1612,99 |
| 1304,65 | 1281,01 | 1332,91 | 1464,66 | 1742,39 | 1614,96 |
| 651,90 | 605,06 | 757,74 | 688,57 | 710,65 | 669,92 |
| 326,41 | 410,77 | 431,96 | 500,24 | 617,04 | 637,09 |
| 408,57 | 422,91 | 466,54 | 488,15 | 563,10 | 619,44 |
| 857,50 | 892,86 | 937,38 | 1074,41 | 1065,86 | |
| 875,89 | 1041,49 | 974,87 | 1559,80 | 1326,90 | 1773,74 |
| 185,63 | 192,29 | 213,69 | 219,49 | 314,37 | 320,61 |
| 197,28 | 217,64 | 250,53 | 199,50 | 286,11 | 262,42 |
| 386,26 | 464,88 | 513,49 | 638,77 | 514,39 | 578,65 |
| 136,36 | 161,81 | 200,93 | 214,24 | 228,01 | 216,26 |
| 673,99 | 725,27 | 644,84 | 585,23 | 918,63 | 1035,26 |
| 369,27 | 473,34 | 501,03 | 637,15 | 839,82 | 825,99 |
| 572,41 | 418,51 | 671,53 | 855,05 | 814,50 | 844,37 |

### 6. Datasets for qRT-PCR study

The experiment was repeated 3 times (values are the quotient between Caveolin-1 and GAPDH expression).

Experiment 1

| | A549 line | | | NL20 line | | |
|---|---|---|---|---|---|---|
| Control | 1,257057 | 1,219772 | 1,121098 | 1,115492 | 1,018236 | 0,9908198 |
| TG10 | 1,445014 | 1,582141 | 1,609119 | 1,034146 | 0,9266025 | 1,031247 |

| | | | | | |
|---|---|---|---|---|---|
| Sidak's multiple comparisons test | Mean Diff. | 95% CI of diff. | Significant? | Summary | Adjusted P Value |
| TG10 - Control | | | | | |
| A549 line | 0.3461 | 0.1079 to 0.5844 | Yes | * | 0.0143 |
| NL20 line | -0.04418 | -0.2824 to 0.1941 | No | ns | 0.8008 |

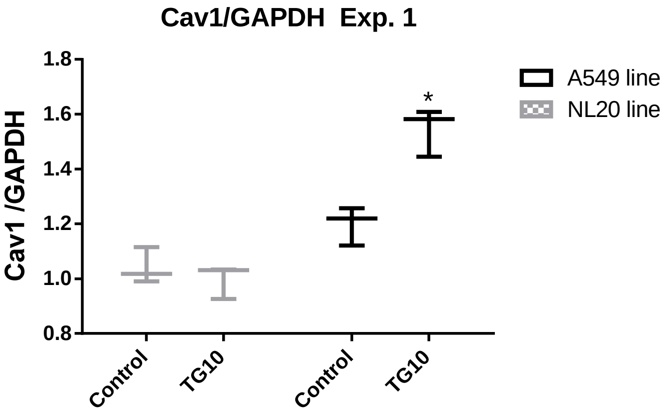

| | | | | | |
|---|---|---|---|---|---|
| Table Analyzed | Cav1/GAPDH 201811 | | | | |
| Two-way RM ANOVA | Matching: Stacked | | | | |
| Alpha | 0.05 | | | | |
| Source of Variation | % of total variation | P value | P value summary | Significant? | |
| Interaction | 19.11 | 0.0157 | * | Yes | |
| tratamiento | 11.44 | 0.0355 | * | Yes | |
| celulas | 62.52 | 0.0005 | *** | Yes | |
| Subjects (matching) | 2.233 | 0.7557 | ns | No | |
| ANOVA table | SS | DF | MS | F (DFn, DFd) | P value |
| Interaction | 0.1143 | 1 | 0.1143 | F (1, 4) = 16.26 | P = 0.0157 |
| tratamiento | 0.06837 | 1 | 0.06837 | F (1, 4) = 9.731 | P = 0.0355 |
| celulas | 0.3737 | 1 | 0.3737 | F (1, 4) = 112.0 | P = 0.0005 |
| Subjects (matching) | 0.01335 | 4 | 0.003337 | F (4, 4) = 0.4749 | P = 0.7557 |
| Residual | 0.02810 | 4 | 0.007026 | | |
| Number of missing values | 0 | | | | |

### Experiment 2

| | A549 line | | | | | | NL20 line | | | | | |
|---|---|---|---|---|---|---|---|---|---|---|---|---|
| Control | 0,91497 | 0,97348 | 1,12069 | 1,23192 | 1,09348 | 1,10249 | 1,00372 | 1,14410 | 1,39692 | 1,25318 | 1,13059 | 1,13079 |
| TG10 | 1,12692 | 1,24071 | 1,35905 | 1,31658 | 1,18018 | 1,17136 | 0,975355 | 1,13423 | 1,23951 | 1,17142 | 1,30580 | 1,14285 |

| | | | | | |
|---|---|---|---|---|---|
| Sidak's multiple comparisons test | Mean Diff. | 95% CI of diff. | Significant? | Summary | Adjusted P Value |
| TG10 - Control | | | | | |
| A549 line | 0.1596 | 0.05158 to 0.2677 | Yes | ** | 0.0061 |
| NL20 line | -0.01502 | -0.1231 to 0.09303 | No | ns | 0.9230 |

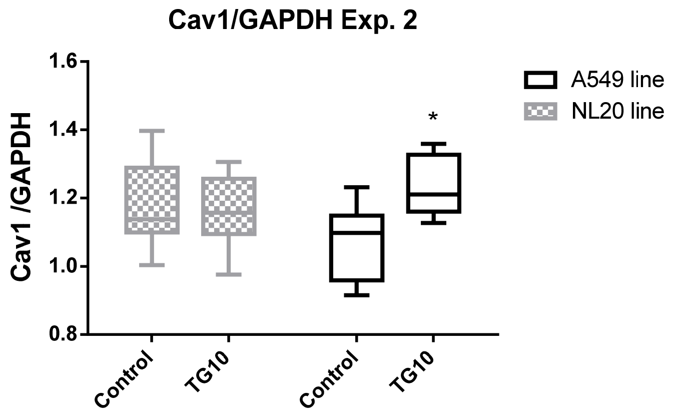

| | | | | | |
|---|---|---|---|---|---|
| Table Analyzed | Cav1/GAPDH 202006 v2 | | | | |
| Two-way RM ANOVA | Matching: Stacked | | | | |
| Alpha | 0.05 | | | | |
| Source of Variation | % of total variation | P value | P value summary | Significant? | |
| Interaction | 13.64 | 0.0133 | * | Yes | |
| tratamiento | 9.349 | 0.0322 | * | Yes | |
| celulas | 0.4802 | 0.7854 | ns | No | |
| Subjects (matching) | 61.40 | 0.0187 | * | Yes | |
| ANOVA table | SS | DF | MS | F (DFn, DFd) | P value |
| Interaction | 0.04575 | 1 | 0.04575 | F (1, 10) = 9.011 | P = 0.0133 |
| tratamiento | 0.03137 | 1 | 0.03137 | F (1, 10) = 6.178 | P = 0.0322 |
| celulas | 0.001611 | 1 | 0.001611 | F (1, 10) = 0.07821 | P = 0.7854 |
| Subjects (matching) | 0.2060 | 10 | 0.02060 | F (10, 10) = 4.057 | P = 0.0187 |
| Residual | 0.05077 | 10 | 0.005077 | | |
| Number of missing values | 0 | | | | |

### Experiment 3

| | A549 line | | | | | | NL20 line | | | | | |
|---|---|---|---|---|---|---|---|---|---|---|---|---|
| Control | 0,2190 | 0,2530 | 0,2810 | 0,2880 | 0,2960 | 0,2670 | 0,2410 | 0,2540 | 0,2760 | 0,2780 | 0,3040 | 0,2810 |
| TG10 | 0,2740 | 0,3020 | 0,3150 | 0,2850 | 0,3130 | 0,2970 | 0,2350 | 0,2630 | 0,2940 | 0,2670 | 0,2850 | 0,2700 |

| | | | | | |
|---|---|---|---|---|---|
| Sidak's multiple comparisons test | Mean Diff. | 95% CI of diff. | Significant? | Summary | Adjusted P Value |
| TG10 - Control | | | | | |
| A549 line | 0.03033 | 0.01104 to 0.04962 | Yes | ** | 0.0041 |
| NL20 line | -0.003333 | -0.02262 to 0.01596 | No | ns | 0.8842 |

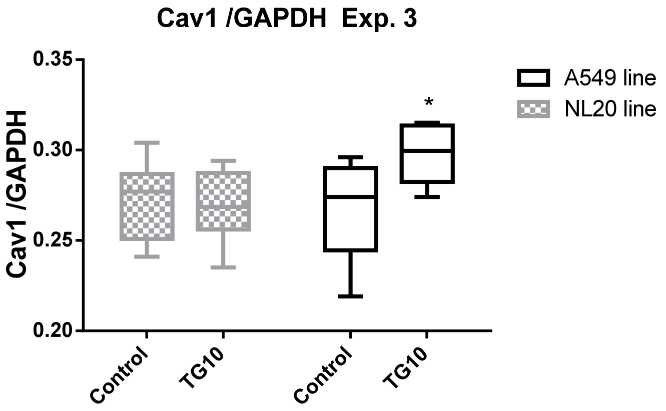

| | | | | | |
|---|---|---|---|---|---|
| Table Analyzed | Cav1/GAPDH 20201104 v3 | | | | |
| Two-way RM ANOVA | Matching: Stacked | | | | |
| Alpha | 0.05 | | | | |
| Source of Variation | % of total variation | P value | P value summary | Significant? | |
| Interaction | 12.68 | 0.0089 | ** | Yes | |
| tratamiento | 8.157 | 0.0265 | * | Yes | |
| celulas | 6.267 | 0.3340 | ns | No | |
| Subjects (matching) | 60.82 | 0.0087 | ** | Yes | |
| ANOVA table | SS | DF | MS | F (DFn, DFd) | P value |
| Interaction | 0.001700 | 1 | 0.001700 | F (1, 10) = 10.51 | P = 0.0089 |
| tratamiento | 0.001094 | 1 | 0.001094 | F (1, 10) = 6.757 | P = 0.0265 |
| celulas | 0.0008402 | 1 | 0.0008402 | F (1, 10) = 1.030 | P = 0.3340 |
| Subjects (matching) | 0.008154 | 10 | 0.0008154 | F (10, 10) = 5.038 | P = 0.0087 |
| Residual | 0.001618 | 10 | 0.0001618 | | |
| Number of missing values | 0 | | | | |
